# Domain-specific proteome remodeling defines mouse myelin aging

**DOI:** 10.64898/2026.07.05.736479

**Authors:** Yarden Freund, Michael Schoof, Noa Moreno, Florian Kriegel, Heather Shin, Vanessa Waelchli, Achint Kaur, Kevin Abraham, Ido Fogel, Sophia M. Shi, Ning-Sum To, Ian H. Guldner, Tom H. Cheung, Andrew C Yang, Andreas Keller, Tony Wyss-Coray, Tal Iram

## Abstract

Myelin, once regarded as a static insulating structure, is now recognized as a dynamic component of the nervous system, capable of remodeling in response to experience. Its breakdown is linked to cognitive decline and neurodegenerative diseases, often preceding neuronal dysfunction. While the myelin proteome has been studied, the age- and sex-related changes it undergoes remain poorly understood. In this study, we purified myelin proteins from young, middle-aged, and aged male and female mice. Using mass spectrometry-based proteomics, we identified 4,095 unique proteins in males and 3,931 in females, with roughly 30% showing significant age-related changes in both sexes. We find an age-related increase in compact myelin structural proteins, such as MBP, MOBP, and CLDN11, and a selective vulnerability in non-compact myelin cytoskeletal proteins, such as SEPTIN2, SEPTIN8, and OPALIN. Notably, disease-associated proteins previously characterized in single-cell transcriptomics appear at the protein level in aged myelin. To firmly distinguish between proteins derived from oligodendrocytes and other cell types *in vivo* we labeled nascent oligodendrocyte proteins with bio-orthogonal non-canonical amino acid tagging (BONCAT). We discovered a dramatic age-related increase in lysosomal and vesicle-associated proteins, while translation and synaptic proteins decrease in myelin with age in both sexes. This dataset highlights molecular mechanisms underlying the loss of myelin integrity and function with age and provides a novel tool for studying oligodendrocyte-derived nascent proteins *in vivo*.

## Introduction

The myelin sheath, a spiraled, multilayered structure formed by oligodendrocytes (OLs) in the central nervous system (CNS), was once thought to be a passive and static feature, primarily responsible for insulating and enhancing the speed of electrical signal transmission across axons^1^. Recent research has redefined it as an active participant in brain plasticity, highlighting its dynamic structure and ability to adapt in response to environmental changes, novel experiences, and during memory formation^2,3^. Additionally, myelin supports neurons metabolically by supplying essential nutrients and energy substrates, such as lactate and pyruvate^4,5^.

While most of the myelin is compacted, some cytoplasm-filled spaces remain in the inner and outer edges of the wrapped sheath (termed inner and outer tongues) and the paranodal loops at both edges of the internode^6,7^, generating a continuum from the oligodendrocyte cytoplasm to the inner layer of the myelin sheath. These non-compact channels contain microtubules^8^, endoplasmic reticulum, and Golgi outposts supporting transport of vesicles and metabolites to the synapse-like periaxonal space between the inner tongue of the myelin and the axon^9^. Indeed, genetic knockout of CNP, critical for maintaining non-compact channel integrity, leads to axonal degeneration^10,11^. These studies support the notion that non-compact myelin channels are critical for myelin structural integrity, proper cellular transport and axonal support. Whether these channels become impaired in aging and neurodegenerative diseases is an area of active research^12^. A recent study identified complex myelination patterns in which multiple internodes are connected by ‘paranodal bridges’ along highly branched axons^13^. With aging, these distal internodes are the most vulnerable to degeneration^13^.

At the clinic, myelin breakdown and white matter lesions are routinely measured by MRI to assess structural abnormalities associated with aging^14^, cognitive decline^15^, and neurodegenerative diseases^16^. These lesions can serve as early indicators of brain aging, often appearing before neuronal degeneration occurs^17^. Recent studies across different species, including in human Multiple Sclerosis lesions, demonstrate that myelin swellings are highly dynamic, revealing that damaged myelin can remodel in response to specific environmental cues^18^. Yet, our understanding of the molecular mechanisms driving these changes and their impact on neurological and cognitive functions remains limited. Our study begins to address this gap by providing a detailed, comprehensive myelin proteome dataset examining age- and sex-related differences, offering a molecular foundation from which mechanistic hypotheses can be drawn and experimentally tested.

## Results

### Age-dependent changes in the myelin proteome

To characterize age-dependent changes in the myelin proteome, we applied TMT-based mass spectrometry to myelin-enriched fractions from male (3, 16, 29 months) and female (3-4, 13, 27 months) mouse brains (Fig. 1a). Quality control confirmed consistent sample preparation, with highly overlapping protein intensity distributions across age groups (Fig. S1a-f). Indeed, myelin-specific proteins, including PLP, MBP, MOBP, MOG, MAG, and CNP, were highly abundant and consistently detected across age groups in the male and female datasets (Fig. 1b-c), confirming successful myelin enrichment. Following quality control and filtering, a total of 4,095 and 3,931 proteins were quantified and retained for analysis in the male and female datasets, respectively. We searched for cell-type-specific markers and found that across all age and sex groups, mature oligodendrocyte proteins were highly detected, while markers of other cell types were lowly expressed or undetected (Fig. 1d, gray box). Together, these results provide high confidence that the purified myelin fraction is predominantly of oligodendrocyte origin. Principal component analysis (PCA) of these proteins revealed a progressive separation of young, middle-aged, and aged groups in both datasets (Fig. 1e-f, Fig. S1g-h), indicating broad age-dependent remodeling of the myelin proteome.

**Figure 1.**
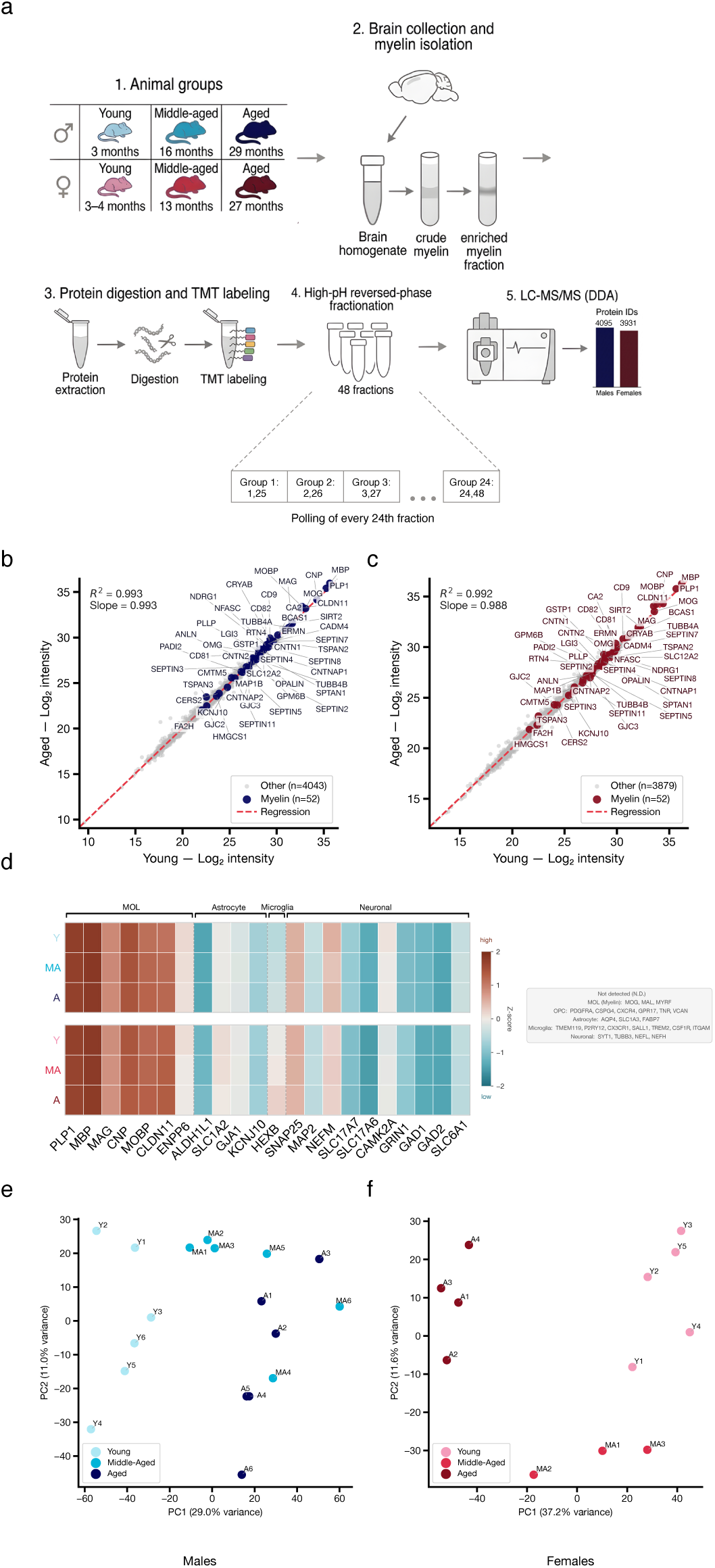
Proteomics workflow and global characterization of the aging myelin proteome in male and female mice. **(a)** Schematic overview of the experimental workflow. Male (3, 16, and 29 months) and female (3–4, 13, and 27 months) mice were grouped by age (young, middle-aged, and aged). Myelin-enriched fractions were isolated from brain homogenates by sucrose gradient centrifugation, followed by protein extraction, tryptic digestion, and TMT labeling. Peptides were pooled and separated into 48 fractions by high-pH reversed-phase fractionation prior to LC-MS/MS analysis (data-dependent acquisition, DDA), yielding 4,095 and 3,931 protein identifications in the male and female datasets, respectively. **(b–c)** Scatter plots of mean log₂ protein intensities in young versus aged mice for males (b) and females (c). Each point represents one protein; myelin-specific proteins (n=38, dark-colored circles) are highlighted against all other quantified proteins (males: n=4,057; females: n=3,893). Known myelin markers (including PLP1, MBP, MOBP, MOG, MAG, and CNP) are labeled and were among the most abundant proteins detected across both datasets. Pearson correlation coefficients (R²=0.993 and R²=0.992, respectively), regression slopes (0.993 and 0.988, respectively), and regression lines were calculated for each scatter plot. **(d)** Heatmap showing z-scored intensities of cell-type-specific marker proteins across young (Y), middle-aged (MA), and aged (A). Markers are shown for MOL, OPC, astrocytes, microglia, and neurons. Proteins not detected (N.D.) are listed in the gray box. **(e-f)** Principal component analysis (PCA) of all quantified proteins in the male (d; n=4,095 proteins) and female (e; n=3,931 proteins) datasets. Each point represents one biological replicate, colored by age group (young, middle-aged, aged).

### Compartment-specific proteomic vulnerability defines myelin aging

We started by analyzing a subset of 56 structural proteins previously validated as present in myelin (Supplementary Table 1)^19,20^, of which 26 showed significant age-related changes in abundance in both sexes revealing sub-compartment specific patterns (Fig. 2a, Fig. S2a, Supplementary Table 2). Compact myelin proteins, including MBP, MOBP, and CLDN11, increased in abundance with age in both sexes (Fig. 2b–d, Fig. S2b-d), which may reflect a compensatory mechanism to preserve myelin compaction. In contrast, non-compact proteins, cytoskeletal and junction components, including OPALIN, GJC3 and several septin family members (including SEPTIN2/5/8) decreased with age (Fig. 2e–g, Fig. S2e-g), potentially underlying altered structural integrity and capacity for cellular transport and axonal metabolic support^21–23^. Together, these findings point to a compartment-specific vulnerability in aging myelin, whereby the non-compacted myelinic channel network is selectively impaired, while compact myelin is maintained. Of note, while total myelin content increases with age^24–28^ (Fig. S2h), in this analysis, we analyze the relative abundance of myelin proteins from an equivalent protein input amount in all age groups. We next sought to place these myelin proteomic changes in the context of oligodendrocyte lineage heterogeneity. Single-cell RNA-seq studies have identified distinct oligodendrocyte subclusters across aging, including disease-associated oligodendrocytes (DAOs) that emerge in aging and neurodegenerative disease models such as Multiple Sclerosis and Alzheimer’s Disease^29–34^ (Fig. 2h–j). Analysis of OL sub-cluster marker proteins defining newly formed, myelinating, and mature oligodendrocyte subtypes (cluster genes adapted from^35^, Supplementary Table 3) revealed distinct age-dependent patterns in the myelin proteome (Fig. 2k–l). Notably, the age-dependent proteomic changes observed in myelin were mirrored at the transcriptomic level, with NFOL, MOL2, and DAO each showing distinct age-associated expression changes that parallel the proteomic trends (Fig. 2h–j). This cross-modal concordance, derived from oligodendrocyte lineage cells isolated specifically from the whole brain, strengthens the conclusion that the observed remodeling of the myelin proteome directly reflects the contributions of distinct oligodendrocyte populations to the myelin sheath during aging.

**Figure 2.**
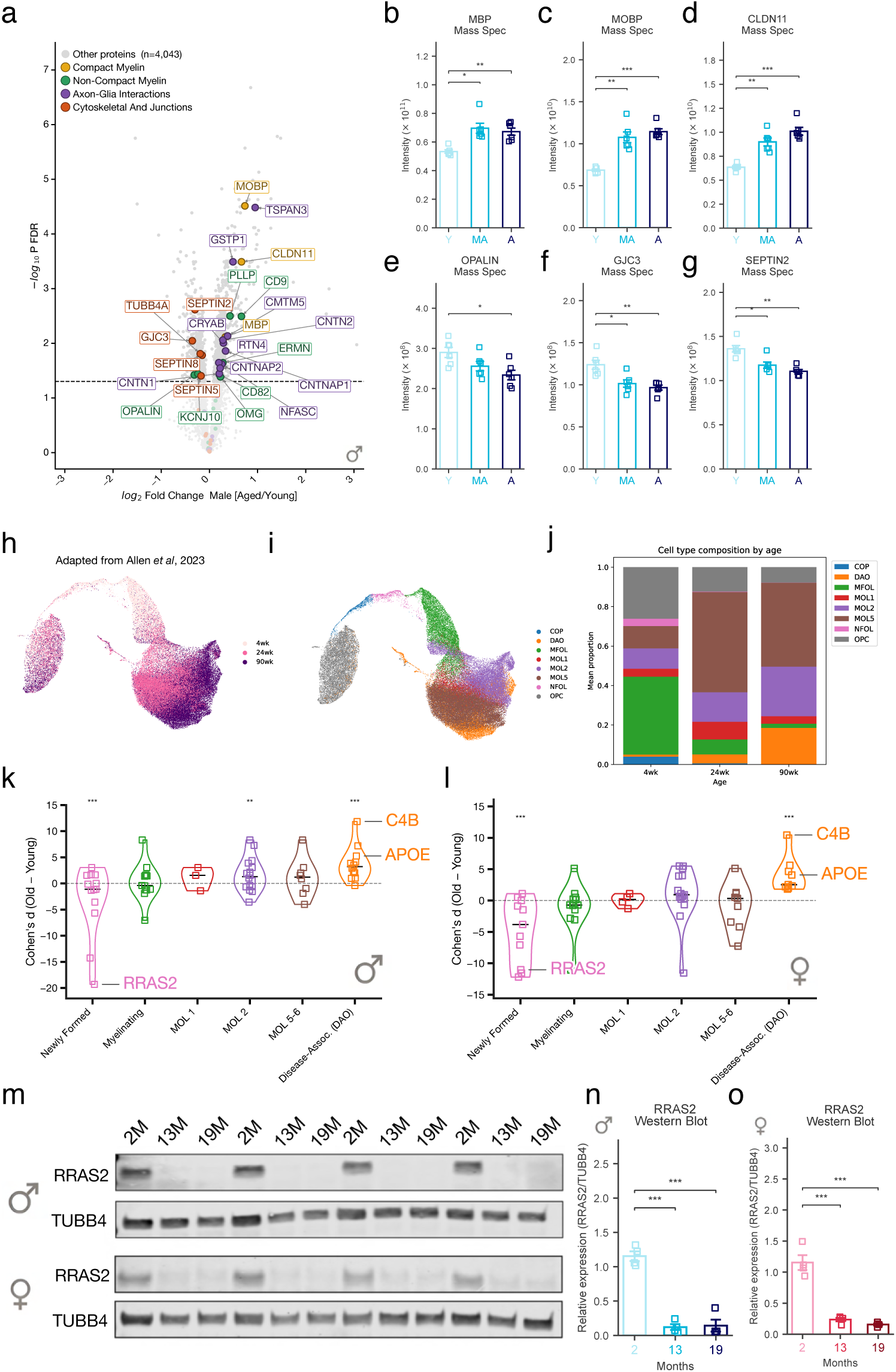
Age-dependent remodeling of the myelin proteome reveals compartment-specific changes and enrichment of disease-associated oligodendrocyte markers. **(a)** Volcano plot showing age-related protein abundance changes (aged versus young) in the male myelin proteome. Proteins are colored by myelin compartment (Compact Myelin, Non-Compact Myelin, Glial Interactions, Cytoskeletal and Junctions; other proteins n=4,037 shown in gray). Significance threshold is indicated by a dashed line (−log₁₀ FDR > 1.3). Significance was assessed by a two-tailed Welch’s t-test with Benjamini–Hochberg correction. Selected proteins are labeled. **(b–g)** Mass spectrometry intensities of individual myelin proteins across age groups (young, middle-aged, aged) in males: MBP (b), MOBP (c), CLDN11 (d), OPALIN (e), GJC3 (f), and SEPTIN2 (g). Statistical pairwise comparisons between age groups were performed using Welch’s t-test with Benjamini–Hochberg FDR correction: *p<0.05, **p<0.01, ***p<0.001. Data are presented as mean ± s.e.m. **(h–j)** Adapted from Allen et al, 2023^35^. UMAP embeddings of oligodendrocyte lineage cells colored by age (h; 4wk, 24wk, 90wk) and oligodendrocyte subtype (i; OPC, COP, NFOL, MFOL, MOL1-6, DAO). (j) Stacked bar chart showing cell type composition by age group (4wk, 24wk, 90wk). **(k–l)** Violin plots showing the distribution of Cohen’s d effect sizes (aged vs. young) for proteins assigned to distinct oligodendrocyte subclusters in males (k) and females (l). Selected proteins (RRAS2, APOE, C4B) are labeled. Permutation-based test, **p<0.01, ***p<0.001. **(m)** Western blot of RRAS2 and TUBB4 in male (top) and female (bottom) brain samples across ages (2M, 13M, 19M). **(n–o)** Quantification of RRAS2 protein levels normalized to TUBB4 by Western blot in males (n) and females (o). One-way ANOVA Kruskal–Wallis test followed by Tukey’s HSD Dunn’s post-hoc pairwise comparisons with Benjamini–Hochberg correction. *p<0.05, **p<0.01, ***p<0.001. Data are presented as mean ± s.e.m. Data are presented as mean ± s.e.m.

Proteins associated with newly formed, actively myelinating oligodendrocytes were relatively decreased in aged myelin (pink violin plot, permutation-based test, p<0.05; Fig. 2k–l), consistent with reduced incorporation of newly generated oligodendrocytes into the myelin compartment^36–39^. This included a decrease in RRAS2, a small GTPase required for oligodendrocyte survival and myelin formation^40,41^, which was further validated by Western blot in an independent aging cohort of both male and female mice (Fig. 2m–o).

In contrast, MOL2-related proteins exhibited a selective upregulation of core structural and metabolic myelin proteins (purple violin plot in Fig. 2k–l), suggesting that a specific mature oligodendrocyte population undergoes targeted remodeling in aging. Notably, markers of disease-associated oligodendrocytes showed significantly greater age-related upregulation in myelin compared to markers from other oligodendrocyte subclusters in both sexes (orange violin plot in Fig. 2k–l). Within the DAO module, complement component 4B (C4B) and Apolipoprotein E (APOE), previously described as disease-associated oligodendrocyte markers at the transcriptomic level^29–34^, are found here at the protein level within purified myelin, suggesting that this stress-responsive OL state gives rise to proteins that are translated and incorporated into the myelin compartment during aging.

Together, these results demonstrate that age-related proteomic changes in myelin arise from compartment-specific remodeling of oligodendrocyte lineage programs, with disproportionate contributions from disease-associated oligodendrocyte subpopulations. The preferential enrichment of disease-associated oligodendrocyte markers in aging myelin indicates that this stress-responsive cellular state not only emerges at the transcriptome^32,42–44^ but also gives rise to proteins that are translated and incorporated into the myelin compartment, suggesting a central role for disease-associated oligodendrocytes in myelin aging and potentially in age-related white matter dysfunction.

### Establishing oligodendrocyte BONCAT lines and characterization of BONCAT proteins in myelin

In order to gain more confidence in the origin of non-myelin-specific proteins in the datasets, we leveraged a tool that enables tagging of cell-type-specific nascent proteomes *in vivo*. BioOrthogonal non-canonical amino acid tagging (BONCAT) enables dynamic proteome tagging by expression of a mutant tRNA synthetase that incorporates azide-bearing non-canonical amino acids (ncAAs) into newly synthesized proteins^45,46^ (Fig. S3a). Using click-based chemistry, tagged proteins are identified by mass spectrometry or located spatially by fluorescent imaging. Building on our previous studies^47,48^, we generated three oligodendrocyte BONCAT lines by crossing Olig2-CRE mice with three BONCAT knock-in mouse lines expressing: (1) mutant phenylalanine tRNA-synthetase flox-stop-flox-eGFP-p2a-PheRS(T413G) (termed Olig2;PheRS*)^48^; (2) mutant tyrosine tRNA-synthetase flox-stop-flox-eGFP-p2a-TyrRS(Y43G) (termed Olig2;TyrRS*)^48^; and mutant methionine tRNA-synthetase (L274G) (termed Olig2;MetRS*)^45,48,49^. In all experiments described, all mice carried a heterozygous allele of the mutant tRNA-synthetase with or without the Olig2-CRE allele. At the age of 8 weeks, all mice received daily i.p. injections of 185 mg/kg of the corresponding ncAAs (AzF, AzY, and Anl, respectively) for five consecutive days (Fig. S3a). Notably, a week before the amino-acid injections, Olig2;MetRS* mice were switched to a low methionine diet as previously described^45,49^. Whole hemisphere protein lysates of all mouse groups were compared by in-gel fluorescence of the azido-modified amino acid (Fig. S3b). Olig2;PheRS* showed superior nascent proteome labeling (red) with similar amounts of total protein loaded in all lanes (green). These results were supported by liquid chromatography with mass spectrometry (LC–MS/MS) analysis of BONCAT-labeled proteins enriched by click-chemistry-based bead pull-down (Fig. S3c-e). After filtering, we detected 2,258 proteins in the Olig2;TyrRS* * line with 77 passing the significance threshold, 3,376 proteins in the Olig2;MetRS* line with 40 passing the significance threshold, and 4,947 proteins in the Olig2-Phe* line with 389 passing the significance threshold (Supplementary Table 4). Lastly, we visualized nascent proteins in Olig2;PheRS* brain sections using fluorescent non-canonical amino acid tagging (FUNCAT) (Fig. S3f-g). Following click-chemistry reaction with Alexa™ 647 Alkyne, nascent proteins (white) were detected exclusively in Cre-expressing (Olig2;PheRS*^+^) animals, with no detectable signal in Cre-negative controls (Olig2;PheRS*^−^). Co-immunostaining with the oligodendrocyte precursor cell (OPC) marker Pdgfrα (green) and the oligodendrocyte transcription factor Olig2 (red) demonstrated that FUNCAT signal co-localized with Olig2-positive cells across the tissue, confirming the cell-type specificity of the labeling (Fig. S3f-g). Together, these data demonstrate that Olig2;PheRS* mice enable specific, efficient, and spatially resolved labeling of the nascent proteome of the oligodendrocyte lineage.

Motivated by this, we next focused specifically on the nascent proteome within the myelin compartment. To this end, we purified myelin from AzF-injected Olig2;PheRS* * mice (Fig. 3a), extracted proteins, and performed BONCAT enrichment. To confirm oligodendrocyte-specific labeling, confocal imaging revealed Alexa™ 647 Alkyne nascent protein signal closely associated with MBP-positive myelin sheaths in Olig2;PheRS* *\** mice (Fig. 3b–c). To our surprise, even when starting the BONCAT enrichment step with a tenth of total protein input material (0.3 mg instead of 3 mg), mass spectrometry analysis of BONCAT-enriched proteins yielded 3,083 unique proteins detected, of which 1,772 were significant above background (p.adj<0.05; Fig. 3d).

**Figure 3.**
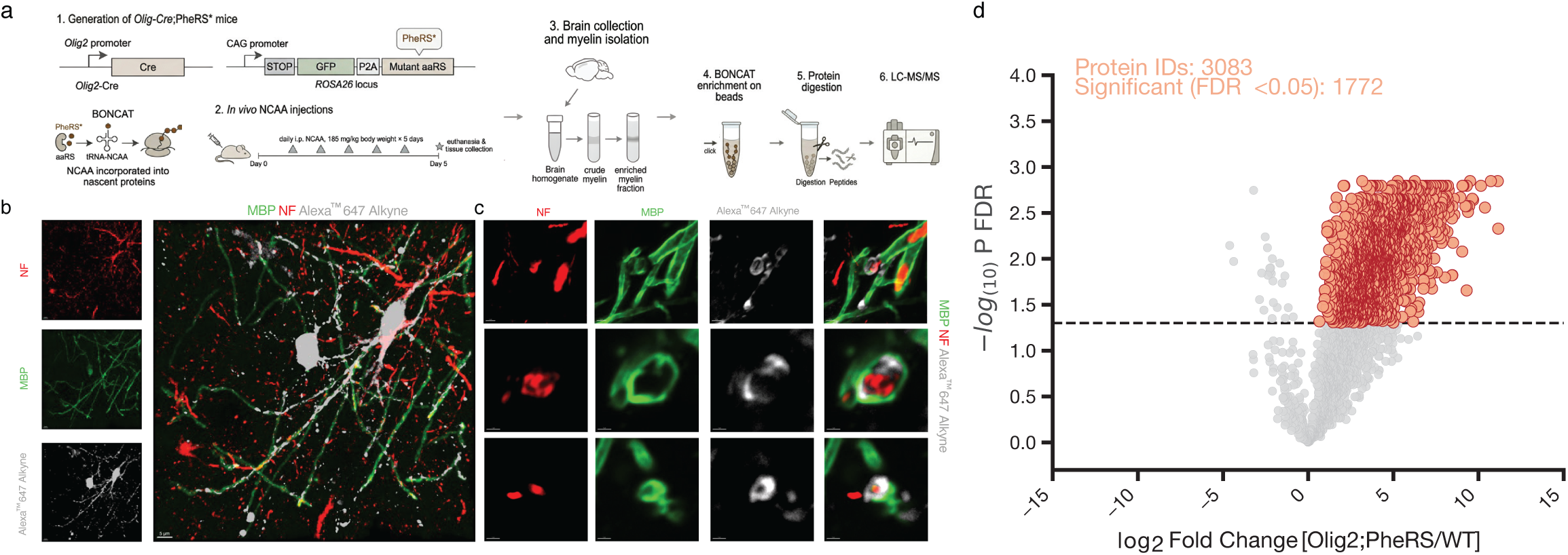
Proteomic characterization of the oligodendrocyte nascent proteome from purified myelin. **(a)** Schematic overview of the experimental workflow. Olig2;PheRS* * mice were generated to enable oligodendrocyte-specific BONCAT labeling of nascent proteins. Mice received daily intraperitoneal injections of the non-canonical amino acid (NCAA; 185 mg/kg body weight) for 5 days, followed by euthanasia and tissue collection. Myelin-enriched fractions were isolated from brain homogenates by sucrose gradient centrifugation, followed by BONCAT enrichment on beads via click chemistry, protein digestion, and LC-MS/MS analysis, yielding 3,083 protein identifications. **(b)** Representative confocal image of a myelinating oligodendrocyte from *Olig2;PheRS\** * mice showing MBP (green), neurofilament (NF, red), and Alexa™ 647 Alkyne nascent protein signal (white), confirming oligodendrocyte-specific BONCAT labeling *in vivo*. Individual channels are shown on the left. Scale bar, 5 µm. **(c)** High-magnification images from *Olig2;PheRS\** * mice showing individual channels (NF, red; MBP, green; Alexa™ 647 Alkyne, white) across three representative cells, illustrating co-localization of the nascent protein synthesis signal with MBP-positive myelin sheaths. Scale bar, 2 µm. **(d)** Volcano plot from LC-MS/MS analysis of BONCAT-enriched proteins from purified myelin of Olig2;PheRS* mice compared to wild-type controls. Dashed line indicates significance threshold (−log₁₀ FDR > 1.3, Benjamini–Hochberg correction). Total protein IDs (3,083) and number of significant hits (1,772) are indicated.

To determine more specifically if neuronal proteins are part of the myelin proteome, we injected a neuronal BONCAT mouse (Camk2a-cre^+/−^;PheRS ^+/–^) for five consecutive days with AzF (Fig.4a). In-gel fluorescence indicated that the tagging worked well (Fig. 4b), yet when we purified myelin and performed BONCAT enrichment, there were only 12 significant neuronal BONCAT-enriched proteins (p-value <0.05 cutoff) (Fig. 4c). These proteins were detected in the total myelin proteome in both the male (Fig. 4d) and female (Fig. 4e) datasets. Further investigation is needed to determine whether these are neuronal proteins that were technically co-purified within the myelin extraction or whether they have a biological function in myelin.

**Figure 4.**
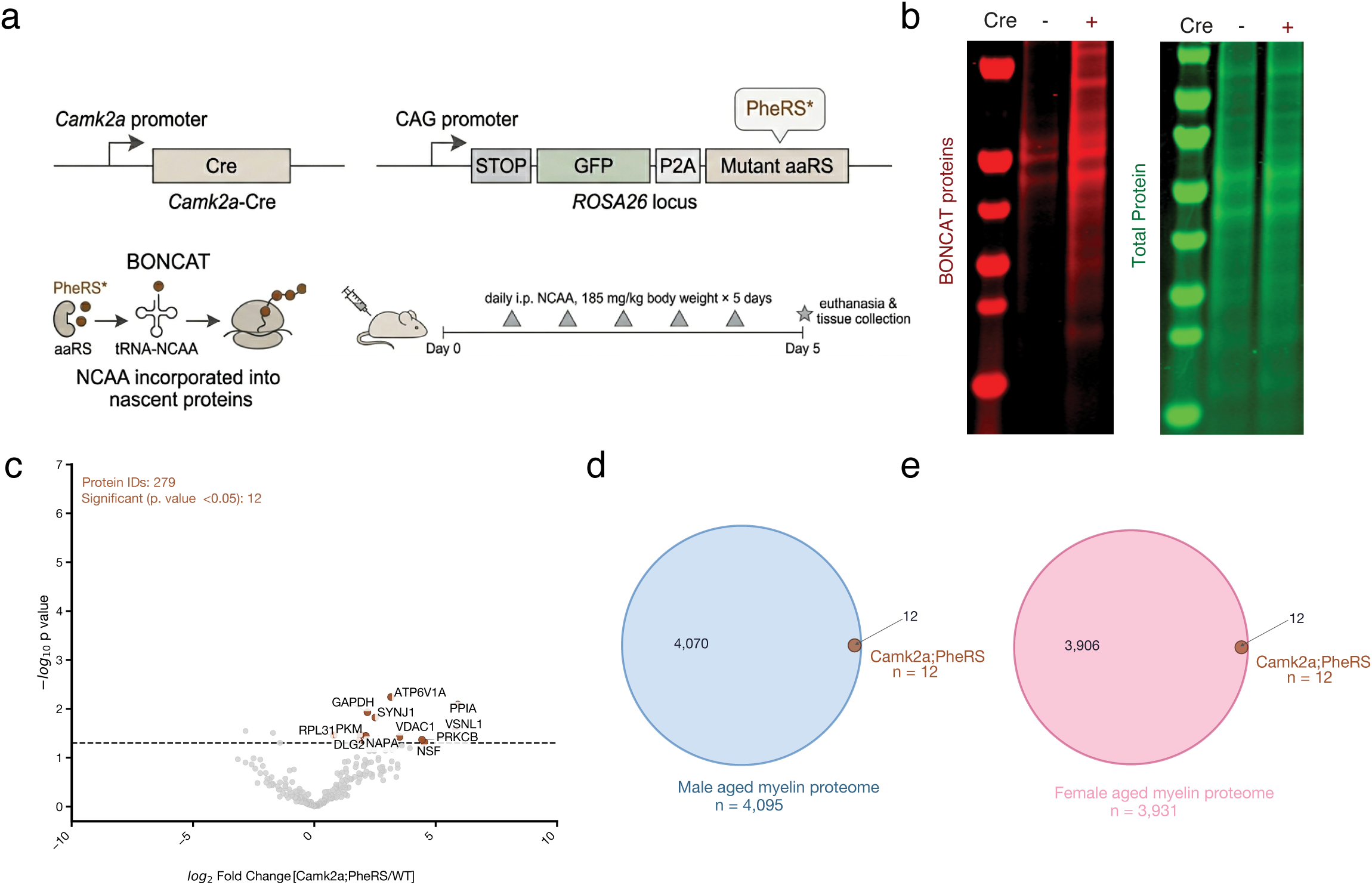
BONCAT-based assessment of neuronal protein contamination in the myelin proteome. **(a)** Schematic of the BONCAT strategy in Camk2a-Cre mice. Camk2a-Cre mice were crossed with PheRS* knock-in mice expressing mutant PheRS* under a CAG promoter at the ROSA26 locus. Mice received daily intraperitoneal injections of AzF (180 mg/kg) for five consecutive days prior to euthanasia and tissue collection. **(b)** In-gel fluorescence of BONCAT-labeled proteins from purified myelin fractions of Camk2a;PheRS*+ (CRE+) and Camk2a;PheRS*− (CRE−) mice. Red channel, azide-modified nascent proteins; green channel, total protein. **(c)** Volcano plot from LC–MS analysis of BONCAT-enriched proteins from purified myelin of Camk2a;PheRS* mice compared to wild-type controls. Dashed line indicates −log₁₀(p-value) < 0.05. Significant proteins are labeled. Total protein IDs and the number of significant hits are indicated. **(d-e)** Venn diagrams showing the overlap between Camk2a;PheRS* myelin BONCAT proteins (n=12, orange) and the total aged myelin proteome from males (D; n=4,095) and females (E; n=3,931).

Overall, this method presents a novel tool for studying nascent and secreted oligodendrocyte proteins across a variety of biological contexts in health and disease.

### Unbiased age-related proteomic alterations in myelin

When comparing the myelin BONCAT proteins with the total aged myelin proteome, 1,465 and 1,437 proteins were shared between the BONCAT dataset and the male and female datasets, respectively (Fig. 5a, e), indicating that a large fraction of BONCAT-identified proteins is represented in the total myelin proteome, supporting their oligodendrocyte origin. We then overlaid myelin BONCAT proteins onto the volcano plots of the male (Fig. 5b) and female (Fig. 5f) total myelin protein datasets to identify age-related changes in the oligodendrocyte-derived fraction. Notably, when examining proteome changes in middle-aged mice (Fig. S4a-d), significant changes among the BONCAT-intersecting proteins were largely confined to the male middle-aged and young comparison (n=272, Fig. S4a), suggesting that myelin proteomic changes occur relatively early during aging, consistent with our previous analysis (Fig. 1d-e). To assess the extent of sex-specific variation, we first examined the correlation between male and female myelin proteomes across age groups, revealing a high degree of similarity (Fig. S5a-c). To further quantify this, we performed Principal Variance Component Analysis (PVCA), which showed that age accounts for a higher portion of the proteome-wide variance, whereas sex contributes minimally (Fig. S5d), indicating that aging, rather than sex, is the predominant driver of proteomic changes in myelin. This finding is consistent with previous reports of limited sex differences in myelin proteomes up to 6 months of age^20^.

**Figure 5.**
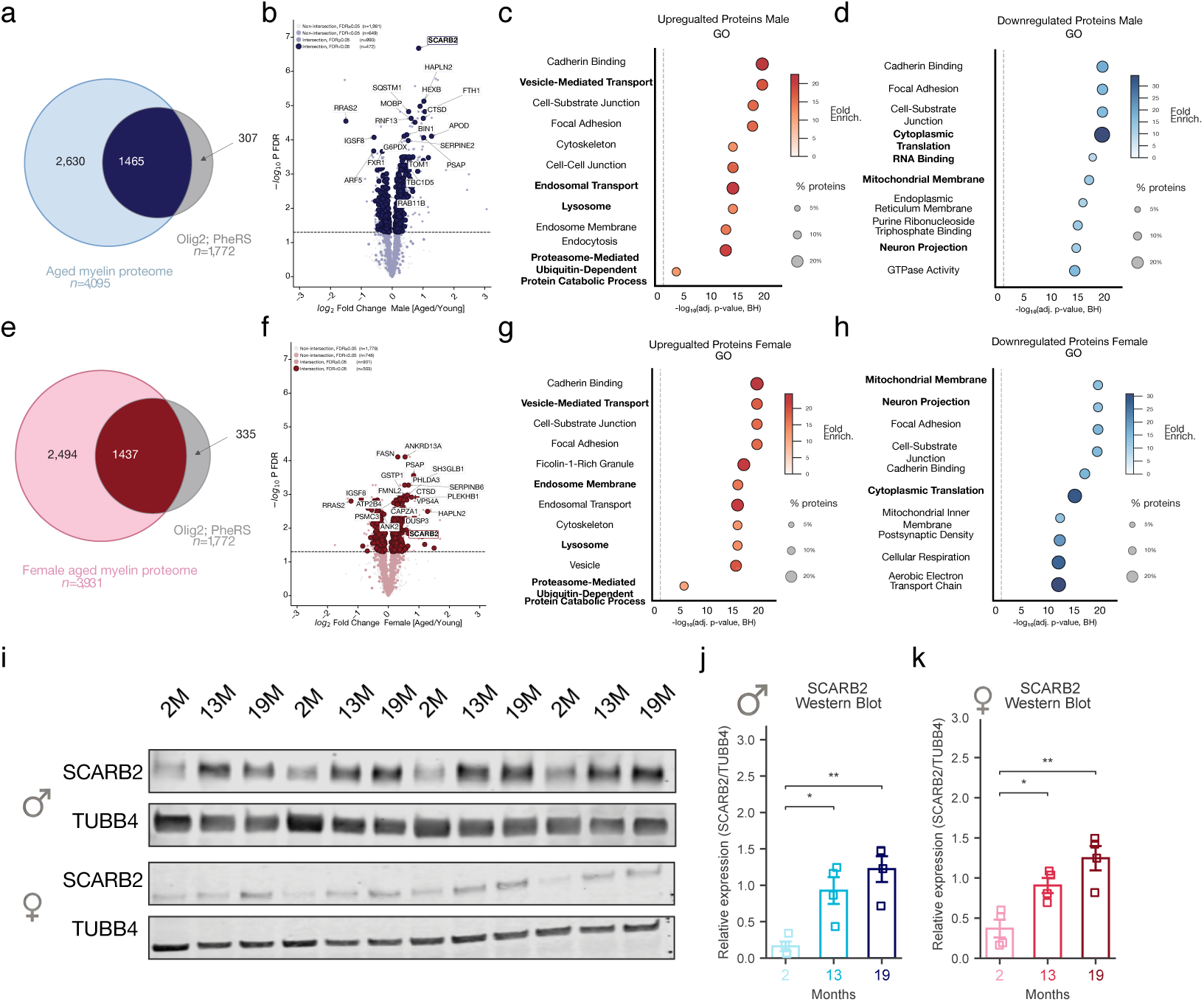
Age-related proteomic alterations in the myelin proteome. (a, **e)** Venn diagrams showing the overlap between the total aged myelin proteome from males (a; n=4,095) and females (e; n=3,931) and myelin BONCAT proteins (n=1,772), yielding 1,465 and 1,437 intersection proteins, respectively. **(b, f)** Volcano plots showing age-related protein abundance changes (aged versus young) in male (b) and female (f) myelin proteomes. Proteins are colored by their overlap with the myelin BONCAT dataset and significance threshold(−log₁₀ FDR > 1.3, dashed line). Intersection proteins passing the significance threshold are highlighted in dark blue (b) and dark red (f), with selected proteins labeled. Significance was assessed by a two-tailed Welch’s t-test with Benjamini–Hochberg correction. **(c, g)** Dot plots showing GO term enrichment analysis (ORA) of proteins significantly upregulated with age in males (c) and females (g). Dot size indicates the percentage of genes per GO term; dot color indicates fold enrichment. Significance is expressed as −log₁₀(adjusted p-value, Benjamini–Hochberg correction). **(d, h)** Dot plots showing GO term enrichment analysis of proteins significantly downregulated with age in males (d) and females (h). Dot size indicates the percentage of genes per GO term; dot color indicates fold enrichment. Significance is expressed as −log₁₀(adjusted p-value, Benjamini–Hochberg correction). **(i)** Representative Western blots of SCARB2 and TUBB4 (loading control) from purified myelin of male (top) and female (bottom) mice across ages (2M, 13M, 19M). **(j,k)** Quantification of SCARB2 protein levels normalized to TUBB4 by Western blot in males (j) and females (k). One-way ANOVA test followed by Tukey’s HSD post-hoc pairwise comparisons with Benjamini–Hochberg correction. *p<0.05. Data are presented as mean± s.e.m.Kruskal–Wallis test followed by Dunn’s post-hoc pairwise comparisons with Benjamini–Hochberg correction. *p<0.05, **p<0.01, ***p<0.001. Data are presented as mean ± s.e.m.

Differentially abundant proteins from the Myelin∩BONCAT intersection in aged versus young myelin were separated into upregulated and downregulated subsets and subjected to overrepresentation analysis against Gene Ontology gene sets to identify age-associated biological processes within the oligodendrocyte-derived myelin fraction. Proteins downregulated with age were enriched for pathways related to translation, cytoskeletal organization, GTPase signaling, and mitochondrial function (Fig. 5d, h). Among the most notable downregulated proteins were multiple components of the cytoplasmic translational machinery, including ribosomal subunits and translation elongation factors, suggesting a broad attenuation of protein synthesis capacity in the aging myelin compartment.

Proteins upregulated with age were enriched for pathways related to cell adhesion, vesicle-mediated transport, and lysosomal and endosomal trafficking in both sexes (Fig. 5c, g), alongside additional proteins associated with lysosomal function and lipid catabolism. Additional upregulated proteins included HAPLN2 and cadherin-associated proteins, suggesting structural remodeling of the myelin-axon interface and the extracellular matrix with age. Notably, while lysosomal proteins are sometimes considered contaminants in biochemically purified myelin, their presence in the myelin BONCAT dataset supports oligodendrocyte production. Among the most significantly upregulated proteins was SCARB2 (LIMP-2), a lysosomal membrane receptor involved in cholesterol export and lipid homeostasis^50^, detected consistently in both males and females (males: log₂FC = 0.856, FDR = 2.1 × 10⁻⁷; females: log₂FC = 0.581, FDR = 0.011). To validate this finding, we selected SCARB2 for Western blot analysis in purified myelin from an independent aging cohort of male and female mice (Fig. 5i). Consistent with the mass spectrometry data, SCARB2 levels progressively increased with age in both sexes, with significant upregulation in middle-aged and aged animals compared with young controls (Fig. 5j–k), confirming the robustness and reproducibility of the age-related proteomic changes identified by mass spectrometry.

## Discussion

This study presents a comprehensive, life-spanning proteomic characterization of myelin in both male and female mice, revealing extensive age-dependent remodeling of the myelin proteome. Across the two sex-specific datasets, we confidently quantified 4,095 and 3,931 proteins in males and females, respectively, spanning three age groups: young, middle-aged, and aged animals, providing a comprehensive view of the myelin proteome. To achieve deep and robust proteome coverage, we employed TMT-based quantitative mass spectrometry coupled with high-pH reversed-phase peptide fractionation, a widely used strategy that reduces sample complexity and enhances peptide identification depth and quantitative precision in complex biological samples^46,51–53^. The resulting data reveal that myelin aging is not a passive process of structural breakdown but rather a complex remodeling characterized by a decline in protein-synthesis capacity, upregulation of lysosomal and endosomal pathways, disproportionate contributions from disease-associated oligodendrocyte subpopulations, and broader changes in cytoskeletal organization and cell adhesion. Furthermore, using BONCAT, we present a novel tool to study nascent oligodendrocyte proteins.

Analysis of a curated set of previously characterized myelin proteins revealed that nearly half showed significant age-related changes in abundance, indicating a substantial reorganization of the myelin compartment with age. It is important to note that these changes represent relative, not absolute, shifts in protein abundance, as equal amounts of myelin protein were loaded across age groups. This is particularly relevant given prior evidence that total myelin content increases with age^24–28^. Compact myelin proteins, including MBP, MOBP, and CLDN11, increased in relative abundance with age in both sexes, which may reflect a compensatory response to preserve myelin compaction integrity or a shift toward a more compaction-dominated myelin proteome, in line with evidence that adult myelin sheaths can undergo structural remodeling in thickness and compaction^54,55^. In contrast, non-compact myelin components showed a selective vulnerability to aging. Cytoskeletal proteins, including septin family members SEPTIN2, SEPTIN5, and SEPTIN8, declined with age, consistent with previous reports implicating septin remodeling in myelin aging^21–23^. Septins are essential for the structural scaffolding of myelinic channels and for supporting axonal metabolic exchange, and their loss may therefore reduce the capacity of myelin to sustain axonal homeostasis^21–23^. Similarly, the gap junction protein GJC3 and the non-compact/paranodal membrane protein OPALIN also declined, suggesting broader disruption of the myelinic channel network and intercellular communication within the myelin sheath^56–59^. Together, these findings point to a compartment-specific vulnerability in the aging myelin proteome, in which the non-compacted myelinic channel network is selectively impaired, while compact myelin is relatively preserved. However, further experimental validation will be required to fully resolve the mechanisms underlying these compositional shifts and their consequences on myelin function.

Age-related proteomic changes in myelin parallel transcriptomic shifts across oligodendrocyte lineage cells^35^, indicating that sheath remodeling reflects broader transcriptional programs in multiple oligodendrocyte populations. With age, proteins associated with newly formed oligodendrocytes decline, consistent with reduced regenerative plasticity, impaired incorporation of new oligodendrocytes, and diminished myelin remodeling capacity. In contrast, markers of the mature MOL2 population are selectively upregulated, including core structural and metabolic myelin proteins, suggesting a shift toward this subtype during aging. Disease-associated oligodendrocyte markers show the strongest age-related increase in myelin in both sexes, raising the possibility of a gain of aberrant or toxic function within the sheath. The selective enrichment of DAO proteins in aging myelin may therefore confer novel, potentially detrimental functions rather than simply reflecting a passive transcriptional state. Among the top candidates are complement component APOE and C4B, both of which have established roles in complement activation and lipid transport, respectively, that are directly relevant to myelin integrity^27–32^. APOE, a major genetic risk factor for late-onset Alzheimer’s disease, is a principal cholesterol and lipid carrier in the brain and plays critical roles in lipid transport, cholesterol metabolism, and myelin repair; its accumulation in aging myelin likely reflects emerging lipid dysregulation with consequences for sheath integrity^60–63^. In the brain, complement proteins, including C1q, C3, and C4, tag synapses and other neural elements for phagocytosis by microglia and macrophages, contributing to complement-mediated synaptic pruning and neurodegeneration^64^. Enrichment of C4b within purified myelin fractions may thus reflect classical complement activity at the myelin surface, as well as potential non-canonical, oligodendrocyte-intrinsic roles. Together, the presence of C4b, APOE, and other DAO markers in purified myelin fractions suggests that a locally pro-inflammatory, metabolically perturbed microenvironment is established within the myelin compartment during aging.

When examining the directionality of age-related changes across the full myelin proteome in an unbiased manner, two opposing and biologically coherent patterns emerged consistently in both sexes. Proteins upregulated with age were enriched for pathways related to lysosomal function, vesicle-mediated transport, and cell adhesion, suggesting a broad activation of the cellular clearance and membrane trafficking machinery in the aging myelin compartment^65^. Among the most prominently upregulated proteins was SCARB2 (LIMP-2), a lysosomal membrane receptor that mediates cholesterol export and lipid homeostasis^50^, whose progressive age-related increase was independently confirmed by Western blot in a separate cohort of aged male and female mice. The upregulation of SCARB2 and associated lysosomal proteins may reflect either a compensatory response to increased demand for lipid turnover and cholesterol redistribution as myelin undergoes age-related structural changes, or alternatively, a marker of dysregulated lipid trafficking that itself contributes to myelin dysfunction, given the dependence of myelin integrity on precise cholesterol metabolism and oligodendrocyte lipid homeostasis^66–68^. Prior ultrastructural and *in vivo* imaging work has shown that aging and stressed myelin can develop focal swellings that accumulate cytoplasm and organelles^18,69^, supporting our interpretation that intracellular trafficking and lysosomal pathways may become locally concentrated within compromised sheaths. In contrast, proteins downregulated with age were enriched for pathways related to translation, cytoskeletal organization, GTPase signaling, and mitochondrial function, suggesting a broad attenuation of the biosynthetic and metabolic capacity of the aging myelin compartment. Determining whether these shifts represent compensatory mechanisms or signs of declining maintenance capacity will require further myelin-specific perturbations *in vivo*.

A key consideration in interpreting myelin proteomic datasets is the potential for contamination from adjacent cellular compartments. Biochemical purification of myelin membranes relies on buoyant density centrifugation, which, while highly enriching for compact myelin, cannot fully exclude contributions from other light membrane compartments with similar biochemical properties, including lysosomes, axonal membranes, and other organellar fractions. The substantial overlap between the BONCAT-validated oligodendrocyte proteome and the total myelin proteome, combined with the high abundance of mature oligodendrocyte markers and the near-absence of markers from other cell types, provides strong evidence that the proteins detected in our myelin fractions are predominantly of oligodendrocyte origin. To directly assess the degree of axonal contamination, a particular concern given the close physical contact of the myelin sheath and the underlying axon, we performed BONCAT labeling in a neuronal line using Camk2a-Cre; PheRS* mice. Strikingly, myelin purification from these animals yielded only 12 significantly enriched neuronal proteins, which were subsequently flagged as potential axonal contaminants in both the male and female datasets. This result indicates that axonal contamination of the myelin proteome is minimal and unlikely to confound the biological conclusions drawn from the dataset. It is important to note, however, that the BONCAT validation was performed in young adult mice, meaning that proteins upregulated with age may not be captured in the BONCAT reference dataset. This represents an inherent limitation of the cross-age validation approach, and proteins detected exclusively in aged myelin fractions should therefore be interpreted with appropriate caution regarding their precise cellular origin.

Beyond its role as a validation strategy in this study, the oligodendrocyte BONCAT system provides a versatile platform for probing the nascent oligodendrocyte and myelin proteome *in vivo*. This is particularly beneficial for cell types with complex morphologies, such as mature oligodendrocytes and neurons, as it enables selective enrichment of the oligodendrocyte-derived nascent proteome directly from tissue lysates and myelin, without the need for prior cellular isolation. Applied to aging or disease models, oligodendrocyte BONCAT could be used to quantify changes in protein synthesis and turnover within the myelin compartment over time, providing a dynamic readout of biosynthetic activity that complements static proteomic measurements. Such experiments could help determine whether the impaired regenerative plasticity and altered cellular communication inferred from our datasets primarily reflect reduced nascent protein production, shifts in protein stability and degradation, or both. More broadly, the ability to selectively label and enrich oligodendrocyte- and myelin-derived proteins from minimally processed tissue positions this approach as a promising tool for biomarker discovery, including the identification of myelin-or oligodendrocyte-enriched proteins that may be released or shed into accessible biological fluids under pathological conditions.

In summary, this study provides a comprehensive proteomic characterization of myelin aging across the lifespan in both sexes, revealing progressive and complex remodeling of the myelin compartment. By integrating unbiased myelin proteomics with oligodendrocyte-specific BONCAT validation, we establish a high-confidence resource for the field and introduce a broadly applicable tool for studying oligodendrocyte and myelin biology *in vivo*. These findings lay the groundwork for future studies to determine whether the proteomic changes identified here are causal drivers or downstream consequences of age-related dysfunction, and to test whether modulating selected age-related myelin pathways can improve myelin integrity in aging and disease.

## Acknowledgments

We thank all members of the Iram and Wyss-Corray Laboratory for their feedback and support.

## Declaration of AI Use

AI-based tools were used to assist with the preparation of this manuscript, specifically Claude (Anthropic) for rephrasing and editing the manuscript text and programming/code development, and Gemini (Google) for generating panel illustrations. All AI-generated content was reviewed, verified, and revised, and the authors take full responsibility for the integrity and accuracy of the published work.

## Contributions

Y.F., M.S., and T.I. conceptualized the project. Y.F. performed most of the experiments and conducted computational analysis of the data. M.S., assisted by K.A., preformed total myelin processing, running on LC-MS/MS and pre-processing the data. N.M, H.S., V.W., and A.Ka. helped establish BONCAT in Olig2;PheRS* mice. N.M and I.F. ran the Western blots. F.K. performed the transcriptomic and clustering analyses, and A.Ke. provided computational expertise and guidance. S.M.S, N-S.T. and T.H.C. performed the LC-MS/MS runs of the BONCAT samples. I.H.G. provided the Camk2a;PheRS* line and assisted with experiments. I.H.G. and A.C.Y. developed wet-bench methodology related to click chemistry. T.I. and T.W-C. supervised the study. Y.F. and T.I. wrote the original manuscript draft and all authors contributed to revising and editing the manuscript.

## Ethics declarations

## Materials and Methods

### Animals

All experiments were conducted in accordance with the Weizmann Institutional Animal Care and Use Committee (IACUC) and Stanford University Laboratory Animal Care guidelines. For mass spectrometry-based proteomics experiments examining age-related changes, wild-type C57BL/6 mice were obtained from the National Institute on Aging (NIA) and either aged in-house at the Stanford Animal Facility for at least one month to acclimate to the facilities or bred in-house. Experiments were conducted at the following ages: males (3, 16, and 29 months) and females (3-4, 13, and 27 months). Olig2-Cre mice (48) were a kind gift of Prof. David Rowitch (University of Cambridge). PheRS(T413G) and TyrRS(Y43G) BONCAT models were generated in collaboration with The Jackson Laboratory (stocks 033734 and 033735, respectively). All other transgenic mouse lines were obtained from The Jackson Laboratory: B6.Cg-Tg(Camk2a-cre)T29-1Stl/J (stock 005359), C57BL/6-Gt(ROSA)26Sortm1(CAG-GFP,-Mars*L274G)Esm/J (stock 028071). A week before ANL injections, MetRS* mice were switched to 0.35% cysteine (low methionine) chow (Envigo, TD.160659). Experiments used male or female mice. Heterozygous cre lines were bred to homozygous BONCAT lines to generate offspring heterozygous for the cre driver (CRE+) or without it (non-CRE) all heterozygous for the BONCAT transgene. For validation experiments (Western blot and immunofluorescence), C57BL/6 mice were obtained from Harlan at 9 weeks of age and aged at the Veterinary Resources Department of the Weizmann Institute to the following timepoints: males (2, 13, and 19 months) and females (4, 13, and 19 months). All mice at the Weizmann Institute were maintained in groups of five per individually ventilated cage under a 12/10h light/dark cycle with autoclaved wood chip bedding. Animals were provided with irradiated rodent hybrid pellet diet and sterile water *ad libitum*.

### NCAA preparation and administration

All AzAAs, including 4-azido-L-phenylalanine (Vector Laboratories; 1406-5G), N-epsilon-azido-L-lysine hydrochloride (Iris Biotech, HAA1625.0005) and 3-azido-L-tyrosine (Watanabe Chemical Industries, J00560) were prepared as a 12.35 mg ml–1 solution for intraperitoneal injections. To hasten the dissolution of AzF, it was first dissolved in 1 M NaOH at 111 mg ml–1, after which it was brought to 12.35 mg ml–1 with sterile 1× PBS. Immediately before intraperitoneal injection at 185 mg kg–1, aliquots were brought to a neutral pH by the addition of 1 M HCl. NCAAs were injected once daily for 5 consecutive days for both BONCAT lines (CRE+ and CRE- mice), following a previously described *in vivo* BONCAT dosing regimen and, with minor adaptations for our experimental condition^48^.

### Tissue Harvesting and Handling

Mice were weighed and euthanized with Pental (Sodium Pentobarbital, 200 mg/ml IP, up to 0.1ml per 10g BW), then transcardially perfused with cold 1× Phosphate-buffered saline (PBS) using a peristaltic pump set to 10 rpm for 1.5 minutes^70^. Following perfusion, brains were immediately extracted and processed according to experimental requirements: either fixed in 4% paraformaldehyde (PFA) or snap-frozen in tubes on dry ice. For histological analyses, brains were fixed in 4% PFA for 24 hours, washed with 1× PBS, and transferred to 30% sucrose solution for cryoprotection until sectioning. For proteomic analyses, snap-frozen tissue was stored at −80°C until further processing.

### Immunofluorescence staining of tissue sections

Free-floating brain sections were washed three times in PBS (1X) for 5 minutes each at 80–100 RPM, then incubated in blocking solution (3% Normal Donkey Serum, 0.3% Triton X-100 in PBS) for 1 hour at room temperature. Sections were then transferred to primary antibody solution prepared in blocking buffer and incubated overnight at 4°C on a shaker at 80–100 RPM. The following primary antibodies were used: rabbit monoclonal anti-PDGFRα (D1E1E; Cell Signaling Technology, #3174; 1:250), mouse monoclonal anti-Olig2 (clone 211F1.1; MilliporeSigma, MABN50; 1:250), rat monoclonal anti-MBP (clone 12; Abcam, ab7349; 1:500), and rabbit polyclonal anti-Neurofilament 200 (MilliporeSigma, N4142; 1:800). After primary incubation, sections were washed three times in PBS (1X) for 5 minutes each and incubated with the appropriate species-specific secondary antibodies diluted in blocking buffer for 1.5 hours at room temperature, protected from light. Sections were washed a further three times in PBS (1X) for 5 minutes each, then mounted onto positively charged slides and allowed to dry. Slides were coverslipped with Vectashield mounting medium containing DAPI and stored at 4°C.

### Microscopy

Images of mounted free-floating 40 µm-thick tissue were captured on a Zeiss LSM 900 (Zeiss) and Nikon Eclipse TI2 (Nikon Microscopes). When capturing images for comparison, all imaging and post-acquisition processing parameters were kept identical across images.

### Copper-catalyzed click reaction on lysates for in-gel fluorescence

Brain tissue was first homogenized by sonication in a strong lysis buffer (8 M urea, 1% SDS, 100 mM chloroacetamide (CAA), 20 mM iodoacetamide (IAA), 1 M NaCl and 1× protease inhibitor in 1× PBS). Sonication was performed for at least 3 cycles of 10 s of sonication with at least 5-s breaks between sonication cycles at an amplitude of 90% using a probe sonicator. Homogenates were centrifuged for 15 min at >16,000g at 4 °C and frozen at −80 °C until further processing. Samples of 110 ug in 33.3 µl were incubated for 1 h with constant shaking to perform the click reaction: 0.83 µl Alexa Fluor 647 alkyne, triethylammonium salt (Thermo Fisher Scientific, A10278) at 5 mM, 1.04 µl copper(II) sulfate (Millipore Sigma, 451657-10G) at 6.68 mM, 2.087 µl THPTA (Click Chemistry Tools, 1010-500) at 33.3 mM, 4.17 µl aminoguanidine hydrochloride (Millipore Sigma, 396494-25G) at 100 mM, 8.33 µl sodium L-ascorbate (Fisher Scientific, A0539500G) at 100 mM and 33.5 µl PBS. Importantly, 20 mM CuSO4 and 50 mM THPTA were mixed at a 1:2 ratio for 15 min before combining the rest of the click reaction. After 1 h of incubation for the click reaction, the reactions were filtered through Zeb Spin Desalting columns, 7 K MWCO, 0.5 ml format (Thermo Fisher Scientific, 89882) following the manufacturer’s protocol to remove unbound fluorophore. The flow-through containing the clicked lysates was retained. Next, 21 µl of the clicked lysates was mixed with 7 µl 1× loading buffer, which was prepared by mixing 10 µl 2-mercaptoethanol (Millipore Sigma, M6250-100ML) with 115 µl 4× NuPAGE LDS sample buffer (Thermo Fisher Scientific, NP0007). Lysate and myelin samples were heated at 95 °C or 40 °C, respectively, for 10 min to denature the proteins. Click-labeled and denatured samples were loaded onto a NuPAGE 12%, Bis-Tris gel (Thermo Fisher Scientific, NP0341BOX) and run at 200 V for 45 min. The gel was imaged to detect the Alexa 647-clicked proteins using a LI-COR Odyssey XF imaging system. To detect total loaded protein, gels were stained with GelCode Blue Stain reagent (Thermo Fisher Scientific, 24590), destained in water for at least 1 h, and then again imaged using a LI-COR Odyssey XF imaging system.

### Copper-catalysed catalyzed click reaction for tissue fluorescence microscopy

After blocking, tissue sections were stained for 2 h in the following click reaction cocktail: 2 µl Alexa Fluor 647alkyne, triethylammonium salt (Thermo Fisher Scientific, A10278) at 5 mM, 5 µl copper(II) sulfate (Millipore Sigma, 451657-10 G) at 20 mM, 10 µl THPTA (Click Chemistry Tools, 1010-500) at 50 mM, 100 µl aminoguanidine hydrochloride (Millipore Sigma, 396494-25G) at 50 mM, 100 µl sodium L-ascorbate (Fisher Scientific, A0539500G) at 50 mM and 783 µl PBS. Importantly, 20 mM CuSO4 and 50 mM THPTA were mixed at a 1:2 ratio for 15 min before combining the rest of the click reaction, as previously described^48^. After staining, tissue sections were washed 3 times in Tris-buffered saline–Tween-20 (TBS-T) and either stained further with antibodies as described in the immunofluorescence staining of tissue sections.

### Myelin Extraction

Myelin was isolated from fresh frozen mouse brains using a sucrose gradient-based ultracentrifugation method for myelin extraction^19^. All solutions were maintained at 4°C throughout the procedure unless otherwise specified. Fresh frozen mouse brains were thawed on ice and dissected in 0.32 M sucrose solution containing cOmplete™ Protease Inhibitor Cocktail, EDTA-Free (Sigma-Aldrich, Cat# 11873580001). Brain tissue was homogenized in a glass tissue grinder. The homogenate was carefully layered over 6 mL of 0.85 M sucrose in ultracentrifuge tubes (Beckman Coulter, 13.5 mL Open-Top Thinwall Ultra-Clear Tube, 16 x 76mm, Cat # 344085), and the remaining volume was filled with 0.32 M sucrose containing protease inhibitors. Paired tubes were balanced to within 0.01 g. Samples were ultracentrifuged at 75,000g for 30 minutes at 4°C in an SW41 Ti swinging bucket rotor (acceleration and deceleration parameters set to 7). The crude myelin fraction, visible as an interphase between the sucrose layers, was carefully collected using a p1000 pipette and diluted in cold distilled water (ddH₂O). This suspension was centrifuged at 75,000g for 15 minutes at 4°C (acceleration and deceleration parameters set to 9) to pellet the myelin. The myelin pellet underwent two consecutive osmotic shock treatments by resuspension in cold ddH₂O, followed by incubation on ice for 10 minutes and centrifugation at 12,000g for 15 minutes. Following the second osmotic shock, the pellet was resuspended in 0.32 M sucrose and subjected to the sucrose gradient ultracentrifugation procedure described above. The final myelin pellet was washed once in ddH2O and centrifuged at 12,000g for 15 minutes. Purified myelin was resuspended in 600 µL of lysis buffer containing 2% sodium dodecyl sulfate (SDS), 1 mM MgCl2, 0.25 U/µl benzonase in PBS, and protease inhibitors in ultra-pure water (UPW) for Mass Spectrometry analysis or Radioimmunoprecipitation Assay buffer (RIPA) and protease inhibitors for Blotting purposes. The suspension was sonicated at 30% amplitude with three pulses (3-second breaks between pulses), repeated 5 times, and then centrifuged at 15,000g for 10 minutes at 4°C. The resulting supernatant was collected and either processed immediately or stored at −80°C for further analysis.

### Myelin protein quantification and statistical analysis

Group differences in myelin protein concentration were assessed using the two-sided Mann–Whitney U test for all pairwise comparisons (young vs. middle-aged, middle-aged vs. aged, and young vs. aged) within each sex, independently and across both sexes combined. No multiple-testing correction was applied to pairwise comparisons within each analysis; the combined sex analysis was included to increase statistical power given the limited per-sex sample sizes (male: n = 6 per group; female: n = 3–5 per group). Data are presented as mean ± s.e.m. with individual data points overlaid. Statistical significance thresholds: *p < 0.05, **p < 0.01, ***p < 0.001. All analyses were performed in Python using the SciPy library (*scipy.stats.mannwhitneyu*).

### Enrichment of BONCAT-labelled proteins and preparation for LC–MS

Brain tissue was first homogenized by sonication in a strong lysis buffer (8 M urea, 1% SDS, 100 mM CAA, 20 mM IAA, 1 M NaCl and 1× protease inhibitor in 1× PBS). The suspension was sonicated at 30% amplitude with three pulses (3-second breaks between pulses), repeated 5 times, and then centrifuged at 15,000g for 10 minutes at 4°C. The resulting supernatant was collected and either processed immediately or stored at −80°C for further analysis, as previously described^46,48^.

Equal protein amounts (3mg of lysate or 300µg for myelin) samples were diluted to 1 ml total with lysis buffer and added to 200 µl dry control agarose beads (Thermo Fisher Scientific, 26150) pre-washed 3 times before sample addition (washed one time with water and two times with 0.8% SDS). Samples were pre-cleared with the control agarose beads to remove nonspecific bead binders by 1 h of light-protected end-over-end rotation. After pre-clearing, samples were centrifuged at 1,000*g* for 5 min to pellet the plain agarose beads. The resultant supernatant was added to 20 µl dry DBCO beads (Vector, 1034-25) pre-washed 4 times before sample addition (washed once with water and 3 times with 0.8% SDS). BONCAT-labelled protein enrichment with the DBCO beads was performed overnight with light-protected end-over-end rotation. After overnight enrichment of BONCAT-labelled proteins to the DBCO agarose beads, 10 µl of 100 mM ANL (Iris Biotech, HAA1625.0005) was added to each sample to quench the DBCO beads to prevent further protein binding. Quenching was performed for 30 min with light-protected end-over-end rotation. After quenching of the DBCO beads, samples were centrifuged for 5 min at 1,000*g* and the supernatant was discarded and the DBCO beads were retained. DBCO beads were washed by the addition of 1 ml water and again centrifuged for 5 min at 1,000*g*. The supernatant was discarded and 0.5 ml 1 mM dithiothreitol (DTT; Thermo Fisher Scientific, R0861) was added to each sample. Samples in 1 mM DTT were incubated for 15 min at 70 °C on a thermomixer set to 70 °C to help to remove proteins nonspecifically bound to the DBCO beads. After incubation, samples were centrifuged for 5 min at 1,000*g* and the resultant supernatant discarded. The DBCO agarose beads were resuspended in 0.5 ml 40 mM IAA (Millipore Sigma, I1149-25G) and incubated light-protected for 30 min to alkylate proteins. After incubation, samples were centrifuged for 5 min at 1000*g* and the resultant supernatant discarded and the DBCO agarose beads were resuspended in 500 µl 0.8% SDS. The DBCO agarose beads were subjected to extensive washing to further remove nonspecifically bound proteins. This was accomplished by washing each sample with 50 ml of 0.8% SDS, 8 M urea and 20% acetonitrile (Fisher Scientific, PI51101). The speed of washes was enhanced by performing them in Poly-Prep chromatography columns (Bio-Rad, 7311550) connected to a vacuum manifold (Fisher Scientific, NC0994627); approximately 7 ml of a wash was added to the column to resuspend the DBCO agarose beads, and then the vacuum was applied to draw through the wash buffer, leaving DBCO agarose beads in the column. Following all washes, DBCO agarose beads were resuspended in 700 µl of 50 mM HEPES (pH 8.0) and immediately transferred to a 1.5 ml tube. DBCO beads were centrifuged for 5 min at 1,000*g*. After centrifugation, the supernatant was completely removed and 200 µl 50 mM HEPES (pH 8.0) (Fisher Scientific, AAJ63002-AE) was added to the DBCO agarose beads. Next, 10 µl of a 0.1 µg µl^−1^ trypsin–Lys-C mix (Promega, V5073) was added to each sample. Proteins bound to the DBCO agarose beads were on-bead digested overnight at 37 °C on a thermomixer set to 1,500–2,000 rpm. The next morning, approximately 16 h after initiating on-bead digestion, samples were centrifuged for 10 min at 1000*g*. Peptides were desalted using Nest Group Inc BioPureSPN Mini, PROTO 300 C18 columns (Fisher Scientific, NC1678001). The desalting process involved conditioning the column with 200 µl methanol for 5 min followed by centrifugation until dry, washing the column twice with 200 µl 50% acetonitrile, 5% formic acid (Thermo Fisher Scientific, 28905) by centrifugation until dry, washing the column 4 times with 5% formic acid by centrifugation until dry, passing peptides through the column by centrifugation, washing the column 4 times wixth 200 µl 5% formic acid, and finally eluting the peptides 2 times with 100 µl 80% acetonitrile, 0.1% formic acid by centrifugation. Following desalting, peptides were dried in a speed vac and then maintained at −80 °C before being run by LC–MS/MS.

### BONCAT LC-MS/MS acquisition

Samples were analyzed using a TimsTOF Pro mass spectrometer (Bruker Daltonics) coupled to a NanoElute system (Bruker Daltonics) with solvent A (0.1% formic acid in water) and solvent B (0.1% formic acid in acetonitrile). Dried peptides were reconstituted with solvent A and injected onto the analytical column, Aurora Ultimate CSI 25 × 75 C18 UHPLC column, using a NanoElute system at 50 °C. For lysate samples (Olig2;PheRS*, TyrRS* and MetRS*), peptides were separated and eluted using the following gradient: 0 min 0% B, 0.5 min 5% B, 27 min 30% B, 27.5 min 95% B, 28 min 95% B, 28.1 min 2% B and 32 min 2% B at a flow rate of 300 nl min–1. For myelin samples of Olig2;PheRS* and Camk2a;PheRS* lysate, peptides were first loaded onto a Waters ACQUITY UPLC M-Class Symmetry C18 Trap Column (100Å, 5µm, 180µm × 20mm) before elution onto the analytical column, using the following gradient: 0 min 0% B; 0.26 min 8% B; 7 min 15% B; 7.1 min 15% B; 27 min 30% B; 28 min 85% B; 28.1 min 0% B until 36.5 min, with a flow rate of 0.45 µL/min from 0–7 min, reduced to 0.3 µL/min from 7.1–35 min, and increased to 0.45 µL/min from 35.1 min until gradient end. All other acquisition parameters were identical for both sample types.

Eluted peptides were measured in DDA-PASEF mode using timsControl 3.0. The source parameters were 1,400 V for capillary voltage, 3.0 lL min^−1^ for dry gas, and 180 °C for dry temperature using Captive Spray (Bruker Daltonics). The MS1 and MS2 spectra were captured from 100 to 1,700 *m/z* in data-dependent parallel accumulation-serial fragmentation (PASEF) mode with 4 PASEF MS/MS frames in 1 complete frame. The ion mobility range (1/K0) was set to 0.85 to 1.30 Vs cm^−2^. The target intensity and intensity threshold were set to 20,000 and 2,500 in MS2 scheduling, with active exclusion activated and set to 0.4 min. 27 eV and 45 eV of collision energies were allocated for 1/K0 = 0.85 Vs cm^−2^ and 1/K0 = 1.30 Vs cm^−2^, respectively.

### LC-MS data analysis

Data captured were processed using Peaks Studio (v.10.6 built on 21 December 2020, Bioinformatics Solution) for sequence database search with the Swiss-Prot Mouse database. Mass error tolerance was set to 20 ppm and 0.05 Da for parent and fragment ions. Carbamidomethylation of cysteine was set as a fixed modification. Protein N-terminal acetylation and methionine oxidation were set as variable modifications, with a maximum of three variable post-translational modifications allowed per peptide. Estimate FDR with decoy fusion was activated. Both FDR for peptides and proteins were set to 1% for filtering.

### Western Blot

Myelin protein samples were retained for protein quantification using a Bicinchoninic Acid (BCA) protein assay kit (Thermo Fisher Scientific, Cat# 23227). Samples were normalized to equal protein concentrations (10 μg total protein) and prepared in 4x Lithium Dodecyl Sulfate (LDS) sample buffer with 10x dithiothreitol (DTT; 500 mM) to achieve final concentrations of 1x LDS and appropriate reducing conditions. Sample volumes were topped up with RIPA buffer. Myelin samples were denatured by heating at 40°C for 5 minutes. Denatured samples were loaded onto pre-cast Bis-Tris gels (Millipore Sigma, Cat# MP41G10) and electrophoresed at a constant voltage of 100V for approximately 90 minutes, until the dye front reached the bottom of the gel. Proteins were transferred to nitrocellulose membranes using a Trans-Blot Turbo transfer system with an appropriate transfer program (7 minutes for low- to medium-molecular-weight proteins). Following transfer, membranes were briefly rinsed three times with distilled water and stained with Ponceau S for 2 minutes to confirm transfer efficiency. Membranes were washed three times with Tris-buffered saline with Tween 20 (TBST; 10 mM Tris-HCl, pH 7.5, 150 mM NaCl, 0.1% Tween-20) for 5 minutes each with gentle shaking. Membranes were blocked with 5% non-fat dry milk or 5% Bovine Serum Albumin (BSA) in TBST for 1 hour at room temperature with gentle shaking. Following blocking, membranes were incubated overnight at 4°C with gentle shaking in primary antibodies diluted 1:1000 in blocking solution. Primary antibodies used were: anti-RRAS (Abcam, Cat# ab154962), anti-beta IV Tubulin (Abcam, Cat# ab179504), anti-LIMPII (Abcam, Cat# ab176317). After primary antibody incubation, membranes were washed three times with TBST for 5 minutes each. Membranes were then incubated with IRDye-conjugated secondary antibodies (LI-COR) diluted 1:10,000 in TBST for 1 hour at room temperature with gentle shaking in light-protected conditions. Following secondary antibody staining, membranes were washed three times with TBST for 5 minutes each. Membranes were imaged using the LI-COR Odyssey imaging system. For loading control analysis, membranes were reprobed with the anti-beta IV Tubulin primary antibody (1:1000) using the same protocol. Western blot validation data were analyzed using one-way ANOVA to assess overall differences between age groups. When the overall ANOVA was significant (p < 0.05), post-hoc pairwise comparisons were performed using Tukey’s HSD test (implemented via the statsmodels package). Statistical significance was set at p < 0.05 for all validation analyses.Western blot validation data were analyzed using non-parametric statistical tests due to the small sample sizes. Kruskal-Wallis tests were performed to assess overall differences between age groups, followed by post-hoc pairwise comparisons using Dunn’s test with Benjamini-Hochberg correction (implemented via the scikit-posthocs package). Statistical significance was set at p < 0.05 for all validation analyses.

### SP2 Protein Clean-up for Protein Purification

Myelin samples were processed using the Single-Pot Solid-Phase-enhanced Sample Preparation (SP2) protocol for quantitative proteomics analysis^71^. Approximately 10 μg of protein per sample was diluted to equal volumes with water to 2% final SDS concentration in 96-well PCR plates based on BCA protein concentration measurements. Sera-Mag Speed Beads A and B (Cytiva, Cat# 45152105050250 and 65152105050250) were mixed 1:1 at room temperature and diluted with four times the volume of water. Beads were washed five times with water using magnetic separation (30 seconds off magnet, 30 seconds on magnet) and stored in 5× volume of water. The SP2 binding solution was prepared, containing absolute ethanol, 15% formic acid, and bead slurry, at a ratio of 15:5:1 μL per sample. Forty microliters of binding solution was added to each sample and incubated for 15 minutes at 500 rpm to ensure complete protein binding. Samples were transferred to filter plates, and unbound fractions were removed by centrifugation at 1000g for 1 minute at room temperature. Beads were washed four times with 200 μL of 70% ethanol by centrifugation at 1000g for 2 minutes, with flow-through discarded after each step.

### Protein Digestion

Trypsin digestion solution was prepared containing 100 mM HEPES, trypsin solution (0.4 μg per sample), 5 mM chloroacetamide, and 1.25 mM Tris(2-carboxyethyl) phosphine hydrochloride (TCEP). Forty microliters of digest solution was added to each cleaned sample on filter plates positioned over collection plates. Plates were sealed with plastic foil and incubated overnight at 37°C, shaking at 500 rpm.

### Peptide Elution and Tandem Mass Tags (TMT) Labeling

Peptides were eluted from beads by centrifugation at 1000g for 1 minute, followed by a second elution with 10 μL of 2% Dimethyl sulfoxide (DMSO) solution. Eluates were dried by vacuum centrifugation for approximately 1 hour. TMT18plex reagents were dissolved in fresh acetonitrile (ACN; 0.5 mg per 68.75 μL), and dried peptides were resuspended in 10 μL water before adding 10 μL TMT reagent according to the experimental labeling scheme. Samples were sealed and incubated for 1 hour at RT, shaking at 500 rpm. The TMT reaction was quenched with 4 μL of 5% hydroxylamine and incubated for 15 minutes at RT, shaking.

### Sample Pooling and Desalting

All TMT-labeled samples corresponding to one experiment were pooled by resuspending each well with 20 μL OASIS Buffer A(0.05% formic acid in LCMS grade water) and combining into a single tube. Wells were washed twice with 180 μL OASIS Buffer A to ensure complete sample recovery. The pooled sample was acidified to pH ∼2 using 15% formic acid to achieve a final concentration of ∼0.5% formic acid. Samples were desalted using Oasis PRiME HLB 96-well µElution Plate, 3 mg Sorbent per Well (Waters, Cat# 186008052). Plates were activated with 100 μL of Oasis B (80% ACN and 0.05% formic acid), washed twice with 100 μL of Oasis A, then loaded with the acidified sample. Following two additional washes with Oasis A, peptides were eluted twice with 50 μL of Oasis B into collection plates, followed by an additional 100 μL wash with the same buffer. Eluates were dried overnight by vacuum centrifugation.

### High-pH Reversed-Phase Fractionation

Dried samples were subjected to high-pH reversed-phase fractionation using Thermo reversed-phase spin columns (Thermo Scientific™, Cat# 84868). Columns were conditioned with 100% Methanol (MeOH), 100% ACN, and 20 mM ammonium formate buffer (pH 10). Samples were dissolved in 250 μL of dilution buffer (20 mM ammonium formate, pH 10) and loaded onto conditioned columns. Following sample loading and washing, peptides were eluted using a step gradient of 2-100% ACN in 20 mM ammonium formate across 48 fractions. These fractions were then pooled into 24 groups for analysis, with every fourth fraction combined (Table 1). Fractions were dried by vacuum centrifugation, resuspended in 0.2% formic acid, and stored at −20°C until Liquid Chromatography-Tandem Mass Spectrometry (LC-MS/MS) analysis.

**Table 1.** High-pH reversed-phase fractionation pooling scheme. High-pH reversed-phase fractionation pooling scheme showing the combination of 48 eluted fractions into 24 final fraction groups. Each grouping row indicates the ACN concentrations (%) used for elution and the corresponding fraction numbers pooled. Every fourth fraction was combined into a single group for downstream LC-MS/MS analysis.

| Groupings | Fraction 1 | Fraction 2 |
| --- | --- | --- |
| Grouping 1 - 2%, 26% | 1 | 25 |
| Grouping 2 - 3%, 27% | 2 | 26 |
| Grouping 3 - 4%, 28% | 3 | 27 |
| Grouping 4 - 5%, 29% | 4 | 28 |
| Grouping 5 - 6%, 30% | 5 | 29 |
| Grouping 6 - 7%, 31% | 6 | 30 |
| Grouping 7 - 8%, 32% | 7 | 31 |
| Grouping 8 - 9%, 33% | 8 | 32 |
| Grouping 9 - 10%, 34% | 9 | 33 |
| Grouping 10 - 11%, 35% | 10 | 34 |
| Grouping 11 - 12%, 36% | 11 | 35 |
| Grouping 12 - 13%, 37% | 12 | 36 |
| Grouping 13 - 14%, 38% | 13 | 37 |
| Grouping 14 - 15%, 39% | 14 | 38 |
| Grouping 15 - 16%, 40% | 15 | 39 |
| Grouping 16 - 17%, 41% | 16 | 40 |
| Grouping 17 - 18%, 42% | 17 | 41 |
| Grouping 18 - 19%, 43% | 18 | 42 |
| Grouping 19 - 20%, 44% | 19 | 43 |
| Grouping 20 - 21%, 45% | 20 | 44 |
| Grouping 21 - 22%, 50% | 21 | 45 |
| Grouping 22 - 23%, 60% | 22 | 46 |
| Grouping 23 - 24%, 80% | 23 | 47 |
| Grouping 24 - 25%, 100% | 24 | 48 |

### TMT-based Mass Spectrometry Data Acquisition and Data Processing

Following high-pH reversed-phase fractionation, peptide fractions were analyzed by liquid chromatography-tandem mass spectrometry (LC-MS/MS). Samples were analyzed by LC-MS/MS using a Dionex UltiMate 3000 RSLCnano System coupled to a Thermo Scientific™ Q Exactive Orbitrap mass spectrometer with Nanospray Flex source. Male and Female myelin samples were fractionated and recombined into 24 total samples for injection. Fractionated samples were reconstituted in 0.1% formic acid in water. Samples were loaded to a trap column (Thermo Scientific™ Acclaim™ PepMap™ C18 column, 2 cm × 100 µm ID, 5 µm) and separated on an analytical column (μPAC™ Neo HPLC Column (110 cm)). Solvent A is 0.1% formic acid in water, and solvent B is 100% ACN with 0.1% formic acid. In a 185-minute method, the gradient was ramped from 5% to 7% at the 10-minute mark, to 28% at the 156.5-minute mark, then to 95% at the 161 to 170-minute mark, before returning to 5% at the 170.1-minute mark for the remainder of the method. The column temperature was set at 50°C. TMT-labeled peptides were analyzed by data-dependent acquisition mode. For the Full MS Scan, the resolution was set to 120,000, the AGC target was 3e6, the Maximum IT was 50 ms, and the scan range was 350 to 1500 m/z. For dd-MS2, Resolution was set to 45,000, AGC target of 1e5, Maximum IT of 120 ms, Loop count of 15, isolation window at 0.7 m/z, fixed first mass of 110 m/z, and NCE of 31.

### TMT Raw Data Processing and Protein Identification

Raw mass spectrometry files were processed using a standardized computational pipeline. Raw files were first converted from .raw format to .mzML format using MSConvertGUI with default parameters and the peakPicking and zeroSamples filters set. Peptide identification and quantification were performed using FragPipe software (v22.0) with the TMT16 workflow (compatible with TMT18plex). Protein identification was performed against the mouse reference proteome database (UniProt, downloaded on May 01, 2025) using MSFragger search engine. Fixed and variable modifications were specified according to the experimental workflow requirements using the FragPipe PTM interface. TMT quantification was performed using FragPipe’s isobaric quantification module at the MS2 level. Protein-level quantification was achieved by grouping peptides to their corresponding proteins. No normalization or log2 transformation was applied during the initial processing, preserving raw abundance relationships for downstream statistical analysis. MaxLFQ algorithm was enabled for additional quantification validation. Results were filtered to include only proteins identified with high confidence, based on a false discovery rate (FDR) of ≤ 1% at both the peptide and protein levels. The FragPipe analysis generated protein abundance tables containing TMT reporter ion intensities for each identified protein across all labeled samples. Additional quality metrics, including peptide spectrum matches (PSMs), sequence coverage, and protein scores, were recorded for each identification.

### Data Preprocessing and Normalization

Myelin proteomics data were processed using a comprehensive normalization pipeline for both male and female datasets. Raw protein abundance data were first filtered to retain proteins with at least 2 PSMs to ensure reliable quantification. Proteins with missing TMT mass channel values were removed from the dataset before normalization. Median normalization was performed by calculating sample-wise medians and normalizing each sample according to the formula:

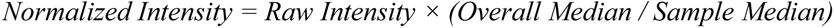

The overall median was calculated across all samples, and the sample median was calculated for each sample. Following normalization, log2 transformation was applied to all intensity values to achieve a normal distribution of the data. Group means were calculated for each age category by sex: males (young: 3 months, Mid: 16 months, Old: 29 months) and females (young: 3-4 months, Mid: 13 months, Old: 27 months) by averaging biological replicates within each group (n = 6 for males, n = 3-5 for females). Z-score normalization was applied row-wise across age groups for trajectory analysis and clustering, where each protein’s expression pattern was standardized to have a mean of 0 and standard deviation of 1 across age groups.

### Quality Control and Normalization Assessment

Comprehensive quality control was implemented to assess the effectiveness of normalization and validate data integrity. Pre- and post-normalization sample median distributions were visualized and compared to ensure successful median centering. Kernel density estimation plots were generated to assess the shape and spread of intensity distributions across samples before and after normalization. Data distribution normality was evaluated using the Shapiro-Wilk test for each age-group means within each sex. Quantile-quantile (Q-Q) plots were generated to visually assess the normality assumption for each group. Despite some deviations from normality, parametric tests were used, given the robustness of t-tests to moderate non-normality and the use of log-transformed data. Within-group variance homogeneity was assessed using multiple statistical measures. For each age group within each sex, the coefficient of variation (CV) was calculated as the ratio of the standard deviation to the mean, expressed as a percentage. Mean and median CV values were calculated across all proteins within each age group to assess technical reproducibility. Sample-to-sample Pearson correlation coefficients were calculated within each age group to evaluate biological similarity. Average correlation coefficients were computed to assess within-group homogeneity. Box plots, violin plots, and correlation heatmaps were generated to visualize patterns of variance and identify potential batch effects or outliers that could compromise statistical comparisons between age groups.

### PCA and Sample Correlation Analysis

PCA and pairwise Pearson correlation analysis were performed independently for male and female datasets on the normalized, log2-transformed protein matrix. Proteins with zero variance were excluded prior to analysis. The remaining missing values were imputed using group-wise medians, and the matrix was z-score-normalized row-wise. PCA was performed using Scanpy, with variance explained by each component indicated on the respective axes. Pairwise Pearson correlation coefficients were computed between all samples and visualized as symmetric heatmaps.

### Statistical Testing

Pairwise comparisons between age groups (young vs. aged, young vs. middle-aged, and middle-aged vs. aged) were performed separately for male and female datasets. Multiple testing correction was applied using the Benjamini–Hochberg false discovery rate (FDR) method, and proteins with FDR < 0.05 were considered statistically significant. Results were visualized as volcano plots displaying log2 fold change on the x-axis and −log10(FDR-adjusted p-value) on the y-axis.

For all BONCAT experiments, statistical comparisons between Cre-positive and Cre-negative animals were performed without a PSM filter. For the BONCAT myelin enrichment experiments (Olig2;PheRS* line), significance was assessed using FDR-adjusted p-values and results were visualized as volcano plots with −log10(FDR) on the y-axis. For the whole hemisphere lysate experiments (Olig2;PheRS* *, Olig2;TyrRS**, and Olig2;MetRS*) and the Camk2a;PheRS* neuronal BONCAT line, results were visualized as volcano plots with −log10(nominal p-value) on the y-axis.

### Single-cell Transcriptomics Analysis of aging Oligodendrocytes

#### Preprocessing and QC

We downloaded the scRNA-seq-imputed spatial gene expression dataset for the aging mouse brain, curated by Allen et al. (2023)^35^ from CellxGene. We subsetted the data to oligodendrocytes (mature + precursors) in whole brain regions. After subsetting, we preprocessed the data using the scanpy package, following best practices (normalization followed by log1p transformation). We confirmed data quality for the already QC-processed data and validated that we obtained 37137 high-quality cells (total_counts > 1000, n_genes_by_counts > 500).

#### Subtype annotation

We ensured consistency in the Transcriptome-Proteome comparison by re-annotating the cells using the same curated marker gene list from the Proteome analysis. For each of the eight Oligodendrocyte subtypes, we computed a cell-type score from the marker gene lists using sc.tl.score_genes, yielding eight scores per cell. We Z-normalized each score and assigned the subtype label corresponding to the highest score for each cell.

#### Visualization

UMAP embeddings for visualization were calculated following best practices: we selected the 3000 most variable genes and computed the neighborhood graph for the UMAP embeddings using the first 50 PCs.

#### Differential Gene Expression Analysis

Count data (including all 7276 genes) were sum-aggregated by donor ID for each mouse in a pseudo-bulk approach. We considered only oligodendrocyte form mice at ages 4 and 90 weeks, resulting in 4 and 5, pseudo-bulk samples, respectively. For Differential Gene Expression Analysis, we employed pyDEseq^72^ (design=”∼ age”) and filtered for significant results (p-value <= 0.05, log2FC > 1).

#### Oligodendrocyte Subcluster Marker Analysis and Violin Plots

Subcluster marker gene lists for mature oligodendrocyte subtypes (NFOL, MFOL, MOL1, MOL2, MOL5–6, and DAO) were curated from published single-cell RNA sequencing studies^29–34^ (Supplementary Table 3). Detected proteins in the myelin proteomics dataset were matched to each subcluster marker list by gene name. For each matched protein, Cohen’s d effect size was calculated for the aged-versus-young comparison, using the pooled standard deviation across biological replicates. The distribution of Cohen’s d values per subcluster was visualized as violin plots with individual data points overlaid. Statistical significance of each subcluster’s mean effect size was assessed using a permutation test (10,000 iterations), in which the observed absolute mean Cohen’s d was compared against a null distribution generated by randomly sampling an equal number of proteins from the full proteome background. Significance thresholds: **p<0.01, ***p<0.001.

### Volcano Plot Overlays

#### Validated myelin proteins

Results were visualized as volcano plots displaying log2 fold change on the x-axis and −log10(FDR-adjusted p-value) on the y-axis. A curated list of 56 previously validated structural myelin proteins was overlaid on each volcano plot and color-coded by myelin compartment category (Compact Myelin, Non-Compact Myelin, Axon-Glia Interactions, and Cytoskeletal and Junctions), regardless of significance. Label positions were optimized using the adjustText algorithm to minimize overlap.

#### Myelin∩BONCAT intersection

Proteins identified in the Myelin∩BONCAT intersection were overlaid onto volcano plots and color-coded based on their membership in the BONCAT-intersecting fraction and significance status (FDR < 0.05). Up to 20 intersection proteins were labeled per plot, with remaining labels selected to maximize spread across the fold-change range. Label positions were optimized using the adjustText algorithm to minimize overlap.

#### Correlation analysis of male and female myelin proteomes

To assess sex-specific variation in the myelin proteome, average protein expression levels were calculated separately for each age group (young, middle-aged, aged) in the male and female datasets. Only proteins detected in both datasets were included. Scatter plots were generated to compare average log2 expression values for males and females in each age group, with Pearson correlation coefficients calculated using scipy.stats.linregress. Proteins were categorized as male-specific, female-specific, or shared based on significant age-related changes (Benjamini-Hochberg-adjusted p < 0.05) identified in the respective young-vs.-aged comparisons.

#### Principal Variance Component Analysis (PVCA)

To quantify the relative contribution of biological variables to proteome-wide variation, Principal Variance Component Analysis (PVCA) was performed on the combined male and female normalized log2 expression matrices. Only proteins detected in both datasets were included (n = 3,639). Missing values were imputed using column-wise median imputation. Principal Component Analysis (PCA) was applied to the scaled expression matrix (samples × proteins) using scikit-learn, retaining sufficient components to explain ≥60% of total variance. Since male and female samples were processed in independent mass spectrometry runs, technical batch effects were accounted for using two complementary approaches. First, batch (mass spectrometry run) was included as a covariate alongside age, sex, and their interaction in the variance decomposition model. Second, batch effects were removed prior to analysis using mean-centering per batch, and PVCA was subsequently performed on the corrected matrix with age, sex, and their interaction as predictors. For each retained PC, an ordinary least squares model was fitted and variance was partitioned using Type-I sum of squares. The proportion explained by each factor was weighted by the corresponding PC’s contribution to total variance, summed across all retained PCs, and normalized to sum to 1, yielding the final estimate of variance attributable to each factor.

#### Pathway Enrichment Analysis

For over-representation analysis, we used the full mouse proteome as the background rather than restricting it to quantified myelin proteins. Because biochemical purification and BONCAT labeling already strongly enrich the measured proteins for myelin and lipid-handling functions, using only quantified myelin proteins as the background would down-weight canonical myelin- and lipid-related pathways and obscure their over-representation within the age-regulated Myelin∩BONCAT fraction. A genome-wide background, therefore, provides a more appropriate reference for identifying biological processes that are selectively emphasized in the age-regulated, oligodendrocyte-derived myelin proteome. Proteins in the Myelin∩BONCAT overlap were then divided into upregulated and downregulated subsets based on the sign of their log2 fold change in aged versus young myelin. Each subset was submitted separately to Enrichr via the gseapy Python package^73^, querying KEGG (KEGG_2019_Mouse), WikiPathways (WikiPathways_2019_Mouse), Gene Ontology Biological Process, Cellular Component, and Molecular Function (GO_2023), Reactome (Reactome_2022), PanglaoDB (PanglaoDB_Augmented_2021), and CellMarker (CellMarker_Augmented_2021). A minimum of three genes was required for enrichment testing, and results were filtered at p.value < 0.05, with an additional Benjamini–Hochberg correction for multiple comparisons. The top 10 significant terms per analysis were visualized as dot plots.

#### Data Visualization and Software

Data visualizations were performed in Python 3.12.5 (Anaconda, Inc.) using the Spyder 5.5.1 IDE or Adobe Illustrator (Adobe, CA, USA).

---

**Supplementary Table 1.** List of 56 structural proteins previously validated to be present in myelin with references.

**Supplementary Table 2.** Raw data and statistics of proteins analyzed in the male dataset (tab 1) and female dataset (tab 2).

**Supplementary Table 3.** Oligodendrocyte subcluster marker proteins defining newly formed, myelinating, and mature oligodendrocyte subtypes.

**Supplementary Table 4.** Raw data of BONCAT-enriched proteins in Olig2;TyrRS* (tab 1), Olig2;MetRS* (tab 2), and Olig2;PheRS* (tab 3) lines in whole hemispherelysates and in myelin from Olig2;PheRS* (tab 4) and Camk2a;PheRS* (tab 5).

