## Supplementary Figures for "Domain-specific proteome remodeling defines mouse myelin aging"

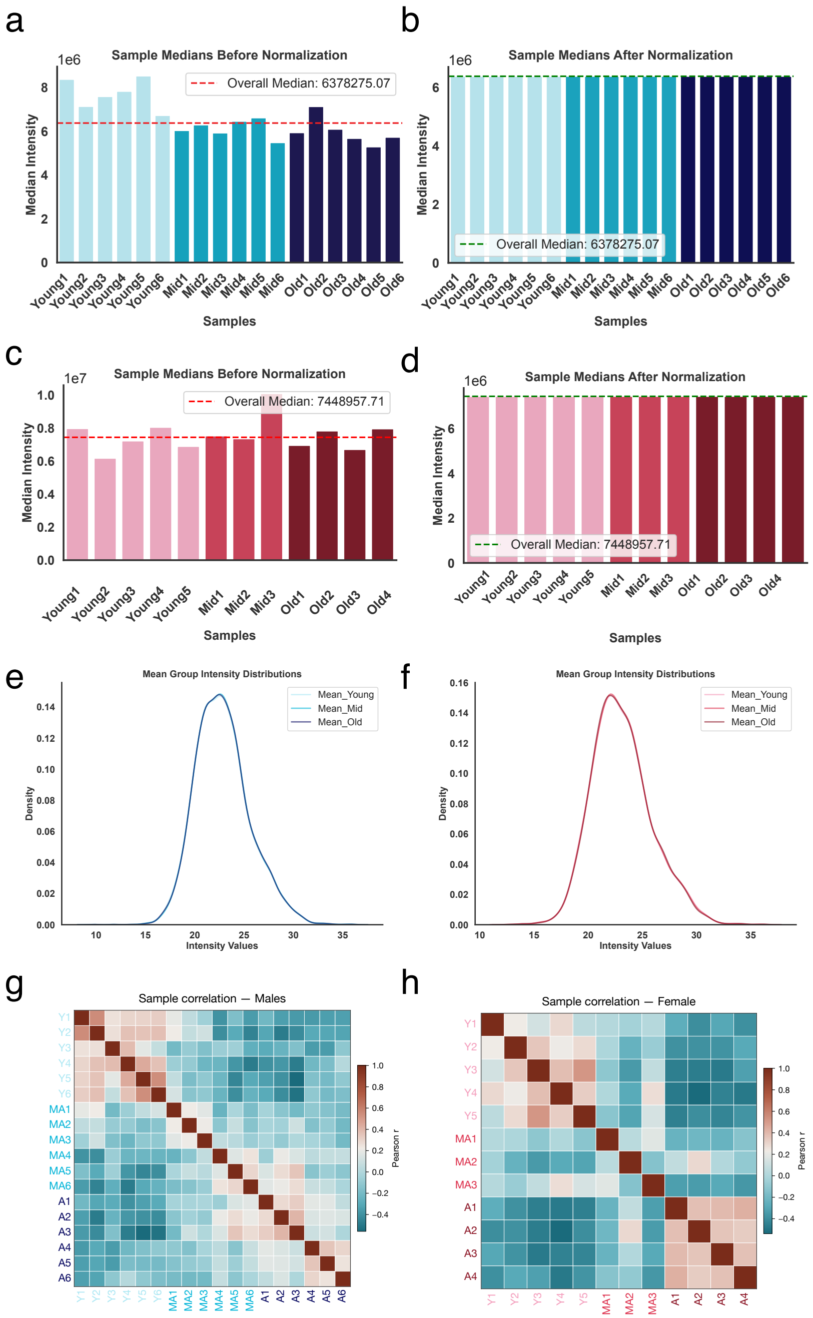


**Supplementary Figure 1**

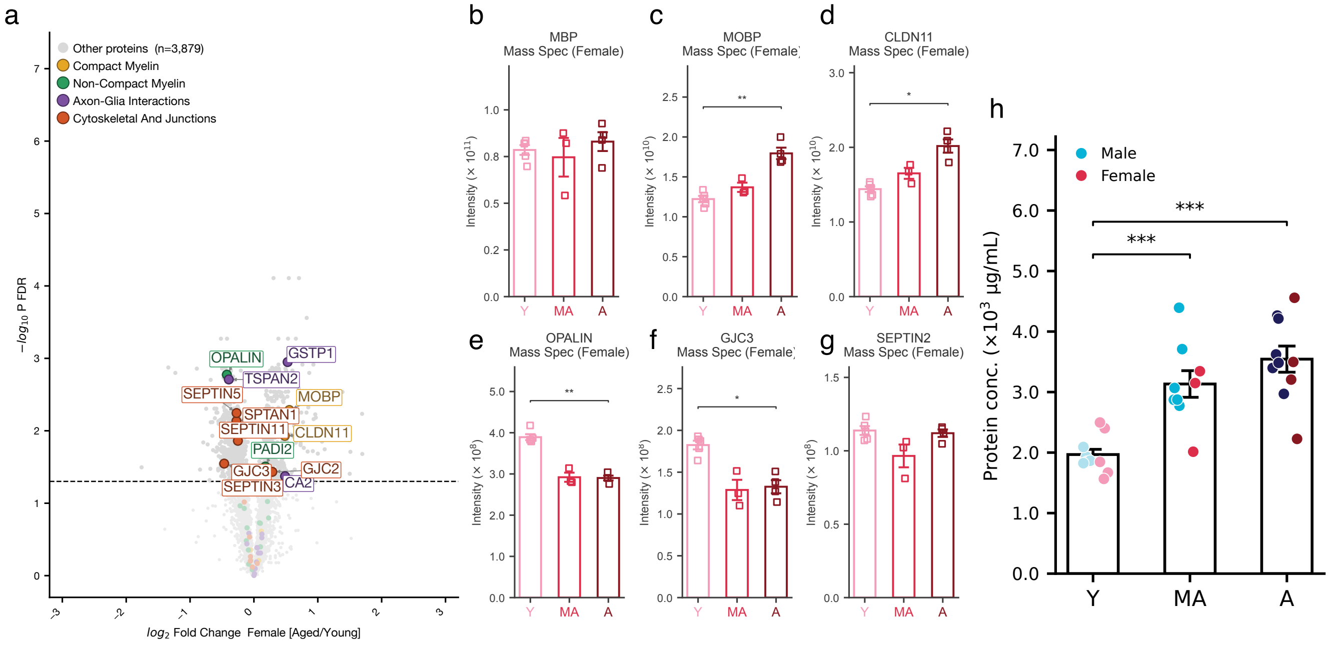


**Supplementary Figure 2**

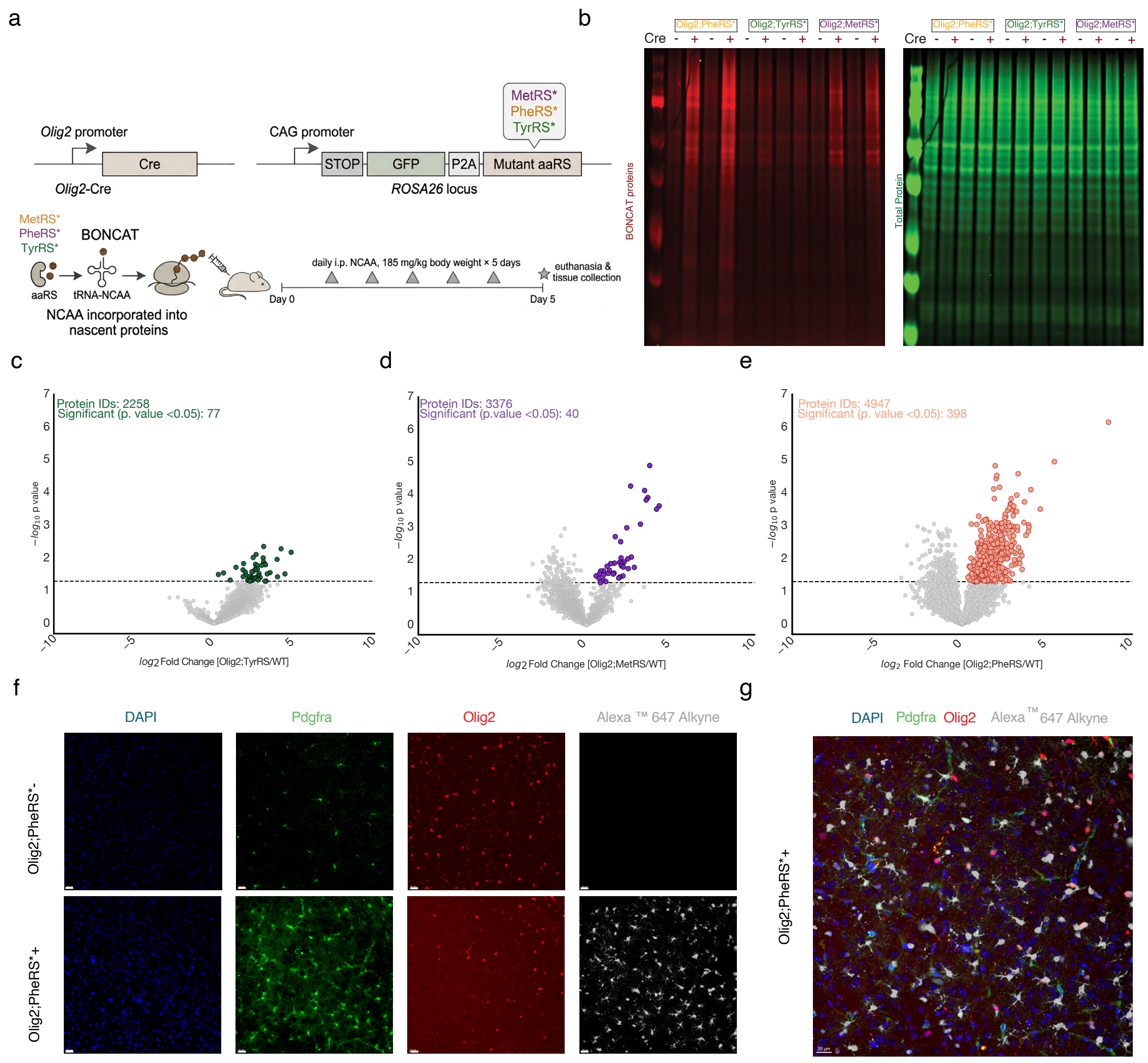


**Supplementary Figure 3**

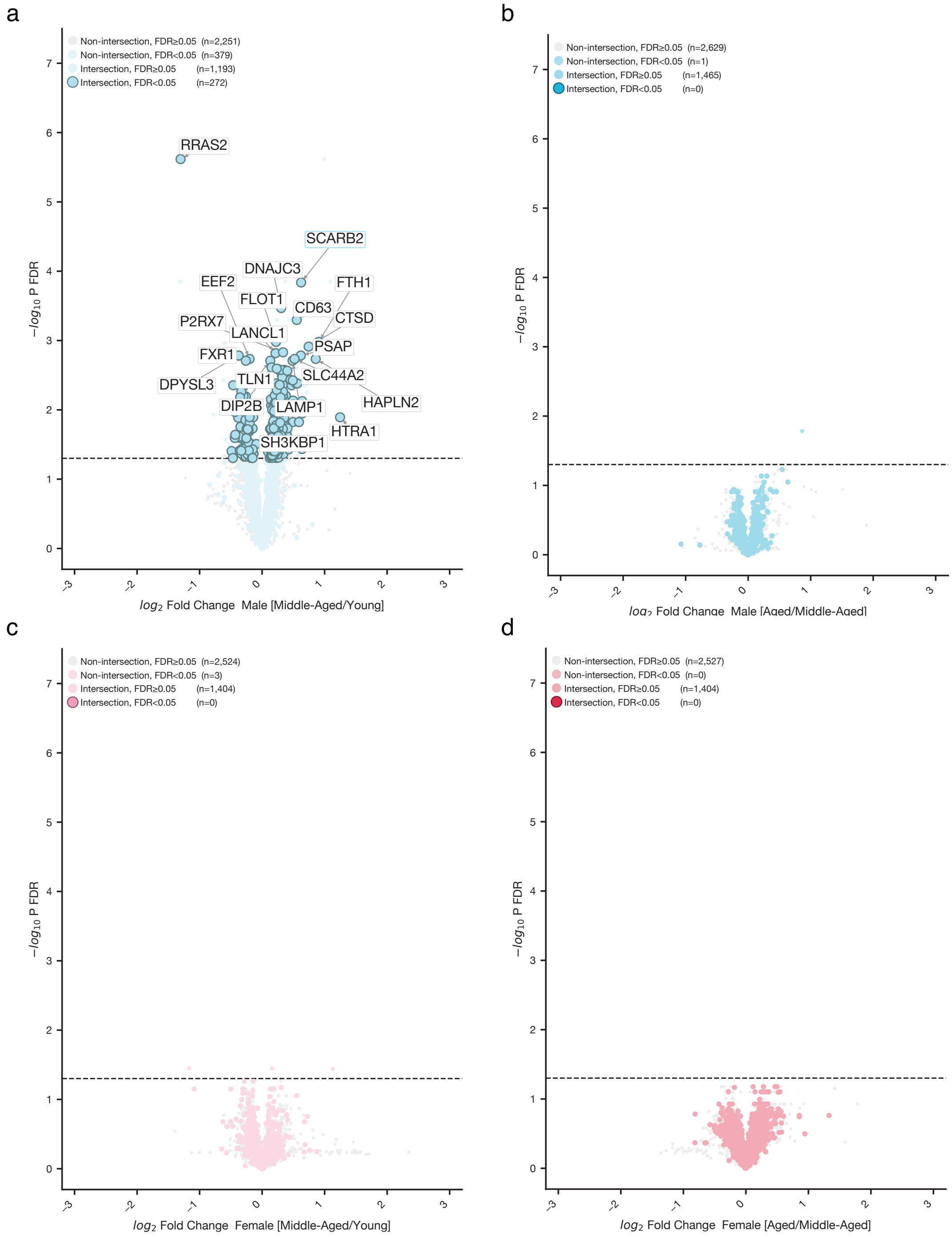


**Supplementary Figure 4**

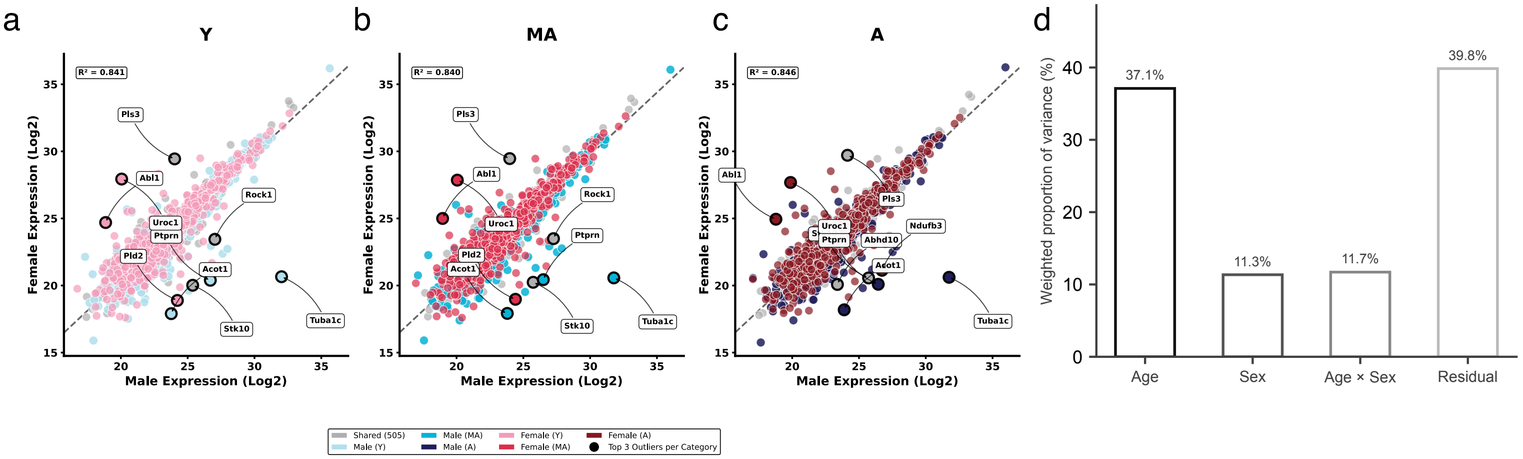


**Supplementary Figure 5**
